# Macropinosomes are a site of HIV-1 entry into primary CD4^+^ T cells

**DOI:** 10.1101/2025.03.07.642068

**Authors:** Tomoyuki Murakami, Ricardo de Souza Cardoso, Praveen Manivannan, Ya-Ting Chang, Eric Rentchler, Kai-Neng Chou, Yipei Tang, Joel A Swanson, Philip D King, Akira Ono

## Abstract

HIV-1 has been observed to enter target cells at both the plasma membrane and endosomes. However, which pathways mediate its entry into primary CD4^+^ T cells, the major targets of this virus, remains unclear. Here, we show that HIV-1 can enter primary CD4^+^ T cells through macropinocytosis, a form of endocytosis. We found that HIV-1 can enter primary CD4^+^ T cells at both the plasma membrane and internal compartments, while entry into common T cell lines occurred primarily at the plasma membrane. Inhibition of macropinocytosis suppressed HIV-1 internalization into and subsequent fusion with primary CD4^+^ T cells regardless of the viral coreceptor usage. Microscopic analysis of viral contents exposed to the cytosol confirmed that HIV-1 fusion occurs at the macropinosomal membrane. Finally, the inhibition of macropinocytosis blocked HIV-1 infection of primary CD4^+^ T cells. Altogether, this study identifies macropinocytosis as one pathway for HIV-1 entry into primary CD4^+^ T cells.

**Significance statement:** HIV-1 entry is an important therapeutic target. However, the exact subcellular location of HIV-1 entry into primary CD4^+^ T cells, a major in vivo host for HIV-1, remains undetermined. The current study shows that macropinosomes serve as a site for productive HIV-1 entry into the cytoplasm of primary CD4+ T cells. Supporting the role for macropinosomes, inhibition of macropinocytosis prevents HIV-1 internalization into, fusion with, and infection of primary CD4^+^ T cells. By contrast, HIV-1 infection of a CD4^+^ T cell line commonly used in HIV-1 research is insensitive to macropinocytosis inhibition. Altogether, this study highlights the primary-T-cell-specific dependence of HIV-1 on macropinocytosis for productive entry and therefore suggests the macropinosome-mediated HIV-1 entry as a potential target for antiviral strategies.

## Introduction

The human immunodeficiency virus type 1 (HIV-1) entry process consists of attachment of a virus particle to target cells and subsequent fusion of viral and target cell membranes. These processes are mediated by the interactions between the HIV-1 envelope glycoprotein (Env) and the receptor CD4 and coreceptors CCR5 or CXCR4 on target cells. While the molecular mechanisms that lead to fusion are well understood, the site of HIV-1 entry remains uncertain.

HIV-1 can fuse with target cells at neutral pH (1, 2), indicating that acidification of endosomal compartments is dispensable for HIV-1 entry. In addition, mutagenesis of the CD4 cytoplasmic tail, which impairs ligand-induced internalization, does not block HIV-1 infection (3, 4). These observations support the possibility that HIV-1 fusion takes place at the plasma membrane. However, using pharmacological or genetic inhibition of endocytosis and live cell imaging, other studies have indicated that HIV-1 enters cells through an endocytic route in a HeLa cell-derived cell line, TZM-bl, and a T cell line, CEM (5). Inhibition of endocytosis either with PitStop2, a clathrin-mediated endocytosis-specific inhibitor, or by expression of dominant negative mutants of dynamin and Eps15 prevents HIV-1 entry into TZM-bl cells (5–7). Furthermore, shRNA screening identified host factors involved in endocytosis as dependency factors for HIV-1 fusion with a CEM-derived cell line (8). However, whether endocytosis plays a role in productive HIV-1 entry into primary CD4^+^ T cells, which are the primary target of HIV-1, is still debated due to the contrasting results obtained with this cell type (9–15).

The membrane-impermeable HIV-1 fusion inhibitor, T-20, completely blocks HIV-1 entry into primary CD4^+^ T cells, even if its addition is delayed after incubation with HIV-1 at 22°C that allows for substantial endocytosis but not fusion (15). This suggests that the plasma membrane is the site of HIV-1 entry into primary CD4^+^ T cells. In contrast to this observation, studies in which the T cells were inoculated at 37°C showed that HIV-1 entry into primary CD4^+^ T cells does occur at a surface inaccessible compartment, e.g., endosomal compartments (13, 14). These studies demonstrated the presence of a virus population that is susceptible to the temperature block at 4°C, which inhibits virus fusion regardless of subcellular location, but is resistant to treatment with fusion inhibitors or protease, which can target only cell-surface fusion events. Consistent with this, a very recent study using live-cell imaging techniques reported that HIV-1 fusion with primary CD4^+^ T cells occurs at neutral pH endosomal compartments (12). Collectively, these findings suggest that HIV-1 can enter primary CD4^+^ T cells via endocytic pathway(s) that are likely to be active at 37°C but not 22°C. However, which endocytic pathway mediates HIV-1 entry into primary CD4^+^ T cells remains to be determined.

There are multiple endocytic pathways, e.g., clathrin-mediated endocytosis, non-clathrin-mediated endocytosis, and macropinocytosis (16). Macropinocytosis is a large-scale and actin-dependent fluid phase form of endocytosis (16, 17). Macropinocytosis is important for sustaining mTORC activity (17–20). It is well known that macrophages, dendritic cells, and cancer cells engage in macropinocytosis (20). Notably, macropinocytosis is implicated in HIV-1 entry into macrophages, endothelial cells, and epithelial cells (21–24). In addition to HIV-1, the entry step of other viruses including ebolavirus, influenza A virus, vaccinia virus, and SARS-CoV-2 is mediated by macropinocytosis (25–29). Recently, our group made the surprising discovery that primary CD4^+^ T cells also engage in macropinocytosis, despite their relative paucity of cytoplasm (17, 19). This, therefore, raises the possibility that HIV-1 could infect primary CD4^+^ T cells through a macropinocytic route. However, whether macropinocytosis in primary CD4^+^ T cells promotes HIV-1 entry into this cell type has not been explored.

In this study, using engineered HIV-1 constructs whose fusion is reversibly arrested, we showed that HIV-1 entry into primary CD4^+^ T cells, but not common T cell lines, can take place at endosomal compartments. We also observed that an inhibitor of macropinocytosis suppressed HIV-1 internalization into and subsequent fusion with primary CD4^+^ T cells. Furthermore, we demonstrated the presence of internalized HIV-1 particles in macropinosomes, defined here as macropinocytosis-derived vesicles, and their fusion at macropinosomes in primary CD4^+^ T cells. Finally, an inhibitor of macropinocytosis prevented HIV-1 infection of primary CD4^+^ T cells. These findings indicate that macropinocytosis enhances HIV-1 infection of primary CD4^+^ T cells via mediating HIV-1 internalization and fusion.

## Results

### HIV-1 entry occurs both at the plasma membrane and endosomal compartments in primary CD4^+^ T cells

To investigate whether productive HIV-1 entry into primary CD4^+^ T cells takes place at endosomal compartments, we examined the sensitivity of HIV-1 encoding the SOS Env protein, which has an engineered disulfide bond between gp120 and gp41 subunits (30–32), to a fusion inhibitor added after inoculation. The fusion process of HIV-1 with SOS Env proceeds up to the interactions of gp120 with CD4 and coreceptors. However, the engineered disulfide bond in SOS Env arrests the subsequent process by preventing gp120 dissociation from gp41. Addition of a reducing agent, DTT, dissociates gp120 and gp41 and thereby allows the remaining fusion steps to occur. Since DTT is membrane permeable, it can induce the fusion of HIV-1 SOS Env regardless of the site of HIV-1 fusion, i.e., both at the plasma membrane and internalized compartments. To distinguish the sites of virus-cell fusion, we used a peptide-based HIV-1 fusion inhibitor, T-20, which inhibits six-helix bundle formation. T-20 can block HIV-1 fusion at the plasma membrane. However, since T-20 is membrane impermeable (33), this drug cannot prevent the fusion at endosomal compartments that are already formed at the time of addition. The combined use of HIV-1 SOS Env, DTT, and T-20 allows us to determine the fraction of HIV-1 entry that occurs at endosomal compartments. An HIV-1 integrase inhibitor, raltegravir (Ral), was used as a positive control because Ral blocks HIV-1 infection regardless of the site of HIV-1 fusion. Activated primary CD4^+^ T cells and two T cell lines frequently used in HIV-1 studies (A3.01 cells and Jurkat cells) were inoculated with CXCR4-using HIV-1_NL4-3/SOS Env_ encoding NanoLuc for 2 hours at 37°C and treated with DTT for 10 minutes at 22∼23°C. At this temperature, the fusion process is arrested at the formation of a pre-hairpin intermediate (34). Cells were then incubated or not with Ral or T-20 for 15 minutes at 22∼23°C and then further cultured for 3 days at 37°C. Additionally, an HIV-1 protease inhibitor, saquinavir (SQV), was added to completely inhibit the unexpected multiple-cycle replication, if any, of the SOS Env-encoding virus during the culture for 3 days. Cells were lysed, and the NanoLuc activity in the lysates was measured and normalized by the total protein amounts (Figure 1A). Although the DTT treatment reduced the total protein amounts by approximately 20% in primary CD4^+^ T cells compared to non-treated cells, neither Ral nor T-20 affected the total protein amounts in DTT-treated cells (Figure S1A). As expected, DTT induced the infection of CXCR4-using HIV-1_NL4-3/SOS Env_ of primary CD4^+^ T cells, which was inhibited by Ral. T-20 was statistically significantly less efficient than Ral in inhibiting HIV-1 infection of primary CD4^+^ T cells isolated from most donors (Figure 1B and S1B), indicating that at least a part of virus-cell fusion takes place at an internal compartment. In contrast, both Ral and T-20 significantly inhibited HIV-1 infection of A3.01 cells and Jurkat cells to a similar extent (Figure 1C and D). These results demonstrate that HIV-1_NL4-3/SOS Env_ entry into primary CD4^+^ T cells can take place at two locations, i.e., at the plasma membrane and internal compartments. In contrast, the plasma membrane is the primary site of HIV-1 entry into A3.01 cells and Jurkat cells.

**Figure 1.**
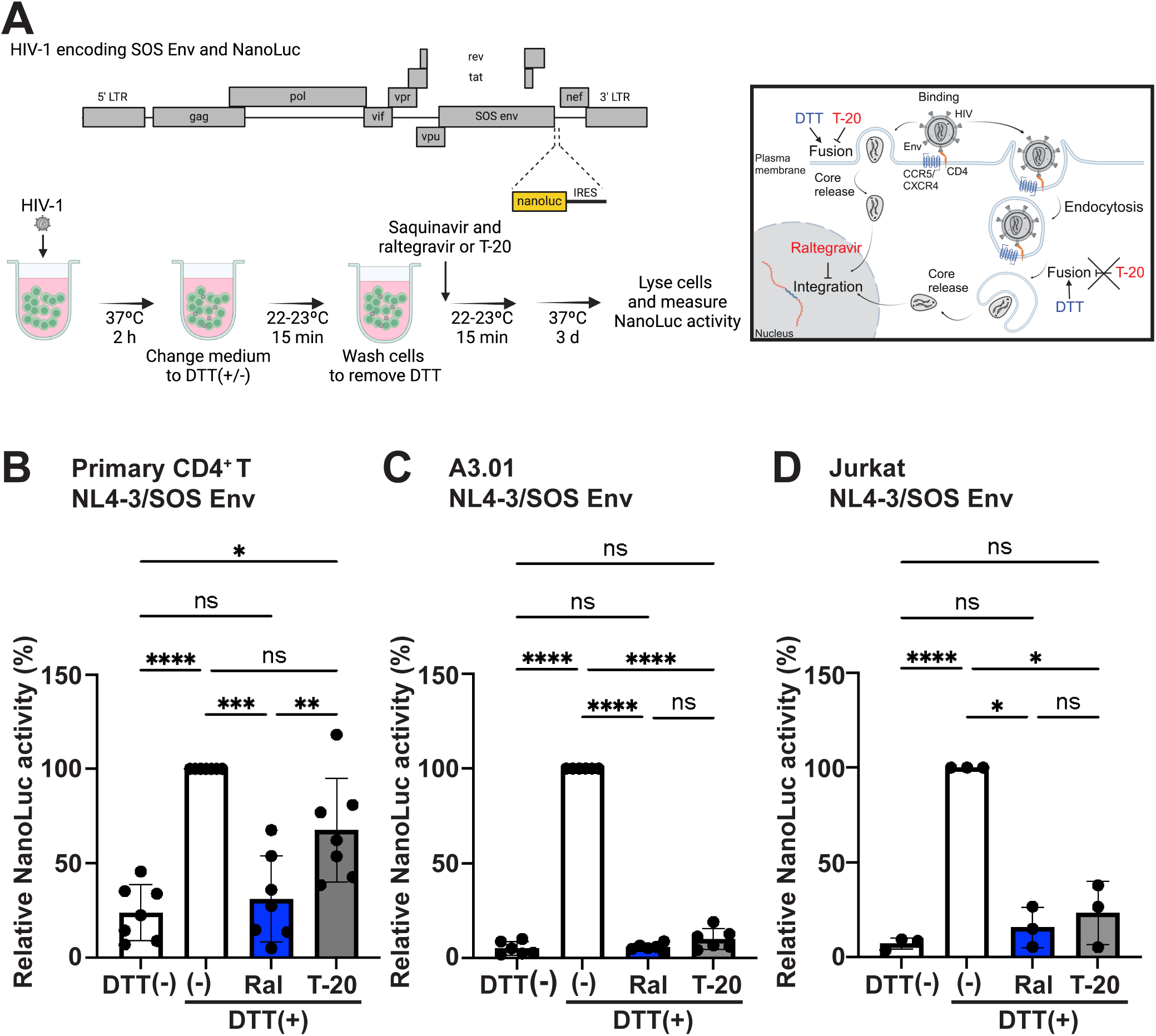
HIV-1 entry into primary CD4+ T cells takes place at both the plasma membrane and internalized compartments. (A) Schematic illustrations of the experimental procedure. Cells were inoculated with HIV-1 molecular clones encoding SOS Env at 37°C for 2 hours. Inoculated cells were incubated for 10 minutes at room temperature (22∼23°C) in the presence and absence of DTT and washed to remove DTT. Cells were then incubated for 15 minutes at room temperature and further cultured at 37°C in the presence of saquinavir and either raltegravir, T-20, or vehicles. After 3 days, the NanoLuc activity in cell lysates was measured and normalized by total protein amounts. (B-D) Activated primary CD4^+^ T cells, A3.01 cells, and Jurkat cells were used as target cells. Indicated cells were inoculated with indicated viral strains. Data are presented as means ±SD. The experiments were performed seven times with primary CD4^+^ T cells isolated from seven donors in B, six times with A3.01 cells in C, and three times with Jurkat cells in D. The *p* values were determined using Tukey’s test following one-way ANOVA. *, *p*<0.05; **, *p*<0.01; ***, *p*<0.001; ****, *p*<0.0001; n.s., not significant.

### Macropinocytosis inhibitors suppress total HIV-1 internalization into primary CD4^+^ T cells

While a previous study showed that endocytosis that is active at 22°C does not mediate HIV-1 entry into primary CD4^+^ T cells (15), our results suggest that endocytosis taking place at 37°C plays a role in HIV-1 infection of primary CD4^+^ T cells (Figure 1). Our group previously demonstrated that primary CD4^+^ T cells engage in macropinocytosis, a fluid-phase endocytosis by which large molecules are preferentially internalized (17, 19). To determine whether macropinocytosis in primary CD4^+^ T cells is active at 22°C, we measured macropinocytosis activity at 4, 22, and 37°C by using a macropinocytosis cargo, BSA. Whereas we observed BSA uptake at 37°C, primary CD4^+^ T cells did not show efficient uptake at 4 and 22°C (Figure S2A and B). This result indicates that macropinocytosis is active at 37°C but not at 4 and 22°C.

To investigate whether macropinocytosis is involved in HIV-1 internalization into primary CD4^+^ T cells, we examined the effect of a macropinocytosis specific inhibitor, EIPA (35). Previously, we observed that 50 μM EIPA shows approximately 90% reduction in macropinocytosis in murine CD4^+^ T cells. By contrast, 10 μM EIPA reduced macropinocytosis activity by only about 10% compared to non-treated cells (19). Treatment of activated human primary CD4+ T cells with 50 μM EIPA for 2 hours and 15 minutes did not affect cell viability (Figure S3A) but significantly reduced macropinocytosis, as measured by the uptake of fluorescent BSA. To examine the effect of EIPA on HIV-1 internalization, we constructed HIV-1 molecular clones encoding the Gag protein with a NanoLuc insertion between MA and CA domains (Gag-iNanoLuc). This virus incorporates NanoLuc into virions. Activated primary CD4^+^ T cells and A3.01 cells were pretreated or not with EIPA or T-20 for 15 minutes and inoculated with the virus for 2 hours at 37°C in the absence or presence of the compounds. Cells were then washed, trypsinized to remove surface-bound viruses for 1 hour at 4°C, and lysed to determine the intracellular NanoLuc activity. Upon HIV-1 fusion, viral contents are released to the cytosol of target cells. Since NanoLuc is a content of virions, the NanoLuc activity in cell lysates should indicate the sum of HIV-1 endocytosis and fusion at the cell surface in this experimental setting. To investigate the contribution of the cell surface-fused HIV-1 to the NanoLuc activity, we used T-20, which inhibits the fusion at the cell surface but not endocytosis of virus particles (Figure 2A). When cells were inoculated at 4°C, at which temperature HIV-1 attaches to the plasma membrane but does not undergo endocytosis and fusion, trypsinization for 1 hour at 4°C removed approximately 90% of HIV-1 from cells compared to those without trypsinization (Figure S4A). This result indicates that trypsin treatment removes almost all surface-bound viruses. When primary CD4^+^ T cells were inoculated with CXCR4-using HIV-1_NL4-3/Gag-iNanoLuc_ at 37°C, both EIPA and T-20 reduced the intracellular NanoLuc activity compared to non-treated cells (Figure 2B). The Nanoluc signal detected in the presence of T-20 is likely due to internalization of fusion-arrested particles via endocytosis. In contrast to primary CD4^+^ T cells, the intracellular NanoLuc activity in A3.01 cells was sensitive to only T-20 and completely insensitive to EIPA (Figure 2C), even though EIPA reduces macropinocytosis in A3.01 cells (Figure S3D). Considering that T-20 is not known to inhibit endocytosis, the reduction in the NanoLuc activity in A3.01 cells is likely due to the inhibitory effect of T-20 on cell surface fusion, thereby increasing the trypsin-accessible virus population. The effect of EIPA on intracellular NanoLuc activity in primary CD4^+^ T cells was not limited to CXCR4-dependent HIV-1. We observed that compared to T-20, EIPA efficiently suppressed intracellular NanoLuc activity of primary CD4^+^ T cells inoculated with CCR5-using HIV-1_NL4-3/Gag-iNanoLuc/CH040env_ in which the env sequence of HIV-1_NL4-3/Gag-iNanoLuc_ was replaced with that of CH040 (Figure 2D). Furthermore, in the same condition, we found that EIPA inhibits internalization of unmodified HIV-1_NL4-3_ into primary CD4^+^ T cells (Figure S4B). EIPA did not affect CD4 density on the surface of primary CD4^+^ T cells (Figure S5A and B), suggesting that the EIPA treatment is unlikely to affect HIV-1 attachment to target cells through CD4. Considering that the macropinocytosis activity measured by BSA uptake is inhibited approximately 40-50% at the concentration of EIPA used in these experiments (Figure S3C), the 30-50% reduction in the intracellular NanoLuc activity in EIPA-treated primary CD4^+^ T cells (Figure 2B and D) is significant. These results strongly suggest that while EIPA-insensitive macropinocytosis and/or non-macropinocytic endocytosis play a role in HIV-1 internalization into both primary CD4^+^ T cells and A3.01 cells, EIPA-sensitive macropinocytosis contributes to HIV-1 internalization specifically into primary CD4^+^ T cells but not A3.01 cells and in a manner independent of the co-receptor usage. To validate the observations obtained with the EIPA treatment, we examined the effect of jasplakinolide in combination with blebbistatin (J/B), which we showed inhibits macropinocytosis in primary CD4^+^ T cells in our previous study (19). We found that J/B treatment does not affect CD4 density on primary CD4^+^ T cells (Figure S5C) but significantly inhibited intracellular NanoLuc activity of primary CD4^+^ T cells inoculated with HIV-1_NL4-3/Gag-iNanoLuc_ (Figure S5D). Altogether, these results suggest that inhibition of macropinocytosis suppresses HIV-1 internalization into primary CD4^+^ T cells.

**Figure 2.**
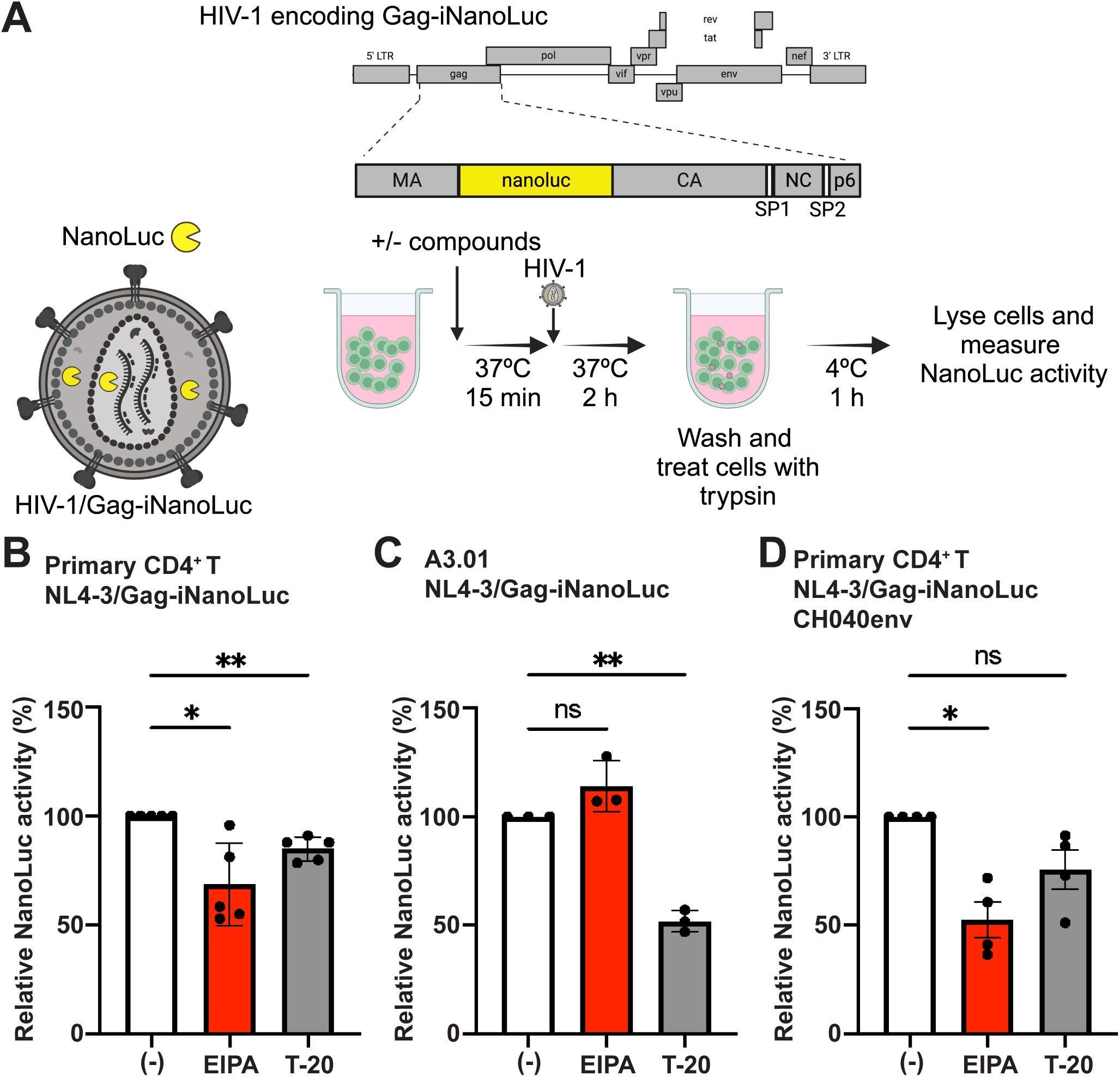
A macropinocytosis inhibitor inhibits total HIV-1 internalization into primary CD4^+^ T cells. (A) A schematic illustration of the experimental procedure performed for panels B-D. Activated primary CD4^+^ T cells and A3.01 cells were pretreated with either EIPA, T-20, or vehicles for 15 minutes at 37°C and inoculated with HIV-1 molecular clones for 2 hours in the presence or absence of either EIPA or T-20. The NanoLuc activity in cell lysates was measured and normalized by total protein amounts. (B-D) Indicated cells were inoculated with indicated viral strains. Data are presented as means ±SD. The experiments were performed with primary CD4^+^ T cells isolated from five (B) and four (D) donors, respectively, in three independent experiments and with A3.01 cells (C) in three independent experiments. The *p* values were determined using Dunnett’s test following one-way ANOVA. *, *p*<0.05; **, *p*<0.01; n.s., not significant.

### Macropinocytosis inhibitors attenuate HIV-1 entry into the cytoplasm of primary CD4^+^ T cells

To test whether EIPA prevents cytoplasmic HIV-1 entry, we performed the BlaM-Vpr-based fusion assay (36), which measures the amount of virion-incorporated BlaM-Vpr released into the cytosol upon HIV-1 fusion with target cells. The β-lactamase activity of BlaM-Vpr in target cells loaded with the cytosolic substrate CCF2 correlates with the efficiency of HIV-1 content release into the cytosol of target cells (Figure 3A). Activated primary CD4^+^ T cells and A3.01 cells were pretreated with the inhibitors, inoculated with HIV-1 containing BlaM-Vpr, loaded with CCF2-AM, and labeled with an anti-CXCR4 or CCR5 antibody. The efficiency of the cytoplasmic entry of HIV-1 (measured as the "fusion" efficiency) and the expression of CXCR4 and CCR5 were analyzed by flow cytometry. We observed approximately 20-25% of reduction in CXCR4 density on the surface of both primary CD4^+^ T cells and A3.01 cells treated with EIPA compared to non-treated cells (Figure 3B-D). Although EIPA suppressed HIV-1_NL4-3_ fusion with both CXCR4^+^ primary CD4^+^ T cells and A3.01 cells, primary CD4^+^ T cells were more sensitive to EIPA than A3.01 cells (approximately 50% reduction versus 25%; Figure 3E-G). To determine whether the reduction in CXCR4 density on the surface of primary CD4^+^ T cells fully explains the suppression of HIV-1_NL4-3_ fusion with CXCR4^+^ primary CD4^+^ T cells, we compared fusion efficiency in non-treated and EIPA-treated cells that expressed the same amounts of CXCR4, as observed among CXCR4 low expressing cells in both populations (Figure S6A and B). In this comparison, EIPA-treated primary CD4^+^ T cells still showed lower HIV-1_NL4-3_ fusion than untreated cells (Figure S6C and D), indicating that the reduction in surface expression of CXCR4 by EIPA does not fully explain the inhibitory effect of EIPA on HIV-1_NL4-3_ fusion with primary CD4^+^ T cells. Consistent with this observation, EIPA also suppressed HIV-1_CH040_ fusion with CCR5^+^ primary CD4^+^ T cells without affecting CCR5 density on the surface of the cells (Figure 3H-J). These results indicate that EIPA attenuates content release of HIV-1 into the cytoplasm of primary CD4^+^ T cells independently of the coreceptor usage. Likewise, J/B inhibited HIV-1_NL4-3_ fusion with primary CD4^+^ T cells without inhibitory effects on CXCR4 expression (Figure S7A and B). Taken together, these results strongly suggest macropinocytosis contributes to cytoplasmic entry of HIV-1 into primary CD4^+^ T cells.

**Figure 3.**
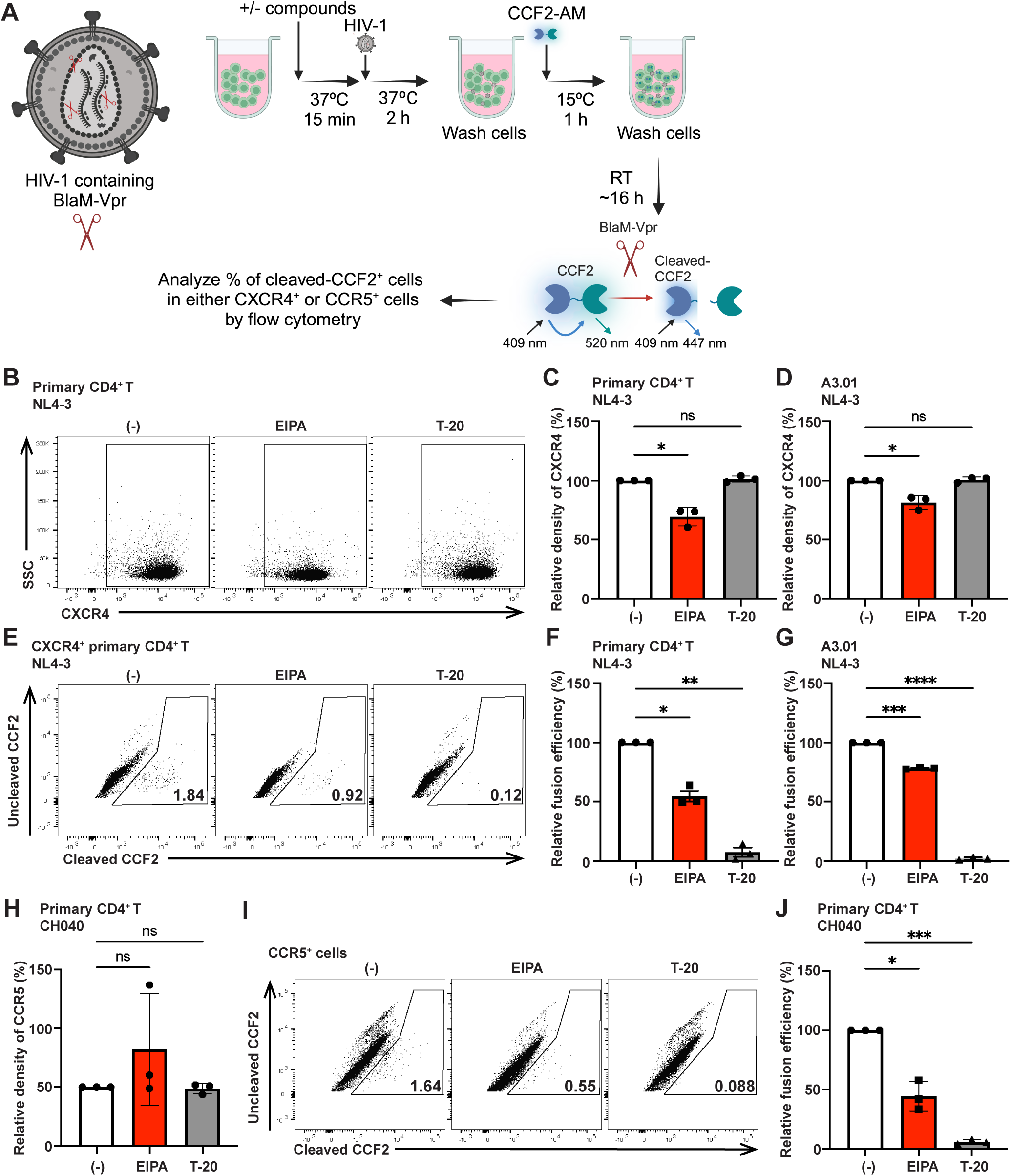
A macropinocytosis inhibitor inhibits HIV-1 fusion with primary CD4^+^ T cells. (A) A schematic illustration of the experimental procedure performed for panels B-J. Activated primary CD4^+^ T cells and A3.01 cells were pretreated with either EIPA or T-20 or left untreated for 15 minutes at 37°C and inoculated with HIV-1 containing BlaM-Vpr for 2 hours at 37°C. Inoculated cells were loaded with CCF2-AM at 15°C for 1 hour and incubated overnight at room temperature. The cells were then labeled with either an anti-CXCR4-APC/Cy7 or CCR5-APC/Cy7 antibody and analyzed by flow cytometry. (B) Representative flow plots of surface expression of CXCR4 on primary CD4^+^ T cells. (C and D) Quantification of the relative density of surface CXCR4 on primary CD4^+^ T cells (C) and A3.01 cells (D). (E) Representative flow plots of the percentages of cleaved CCF2^+^ cells in CXCR4^+^ cells are shown. (F and G) Quantification of the relative percentages of cleaved CCF2^+^ cells (fusion efficiency) in CXCR4^+^ primary CD4^+^ T cells (F) and A3.01 cells (G). (H) Quantification of the relative density of surface CCR5 on primary CD4^+^ T cells. (I) Representative flow plots with the percentages of cleaved CCF2^+^ cells in CCR5^+^ cells are shown. (J) Quantification of the relative percentages of cleaved CCF2^+^ cells (fusion efficiency) in CCR5^+^ primary CD4^+^ T cells. Data are presented as mean ±SD in C, D, F-H, and J. The experiments were performed three times with primary CD4^+^ T cells isolated from three donors in B, C, E, F, and H-J. The experiments were performed three times with A3.01 cells independently in D and G. The *p* values were determined using Dunnett’s test following one-way ANOVA in C, E-G. *, *p*<0.05; **, *p*<0.01; ***, *p*<0.001; ****, *p*<0.0001; n.s., not significant.

### A microscopy analysis displays macropinocytosis-mediated HIV-1 particle internalization into primary CD4^+^ T cells

The enzyme-based bulk cell analyses described above support the involvement of macropinocytosis in HIV-1 internalization into and fusion with primary CD4^+^ T cells (Figures 2 and 3). To test whether HIV-1 particles are internalized through macropinocytosis in primary CD4^+^ T cells, we performed microscopic analyses using HIV-1_NL4-3/Gag-iVenus_ containing mScarlet-Vpr. Labelling of HIV-1 particles with two fluorescent proteins allowed us to distinguish unambiguously HIV-1 particles from CD4^+^ T cell-derived autofluorescent punctate signals, which are either in red or green fluorescence but not both. Activated primary CD4^+^ T cells were pretreated or not with EIPA or T-20 and inoculated with double-labeled HIV-1 in the presence of BSA-AlexaFluor647 for 1 hour at 37°C, washed, trypsinized to remove surface-bound HIV-1 particles, and immunostained with an anti-CD4 antibody conjugated with a fluorescent dye, BV421. Cells were then fixed and observed using a spinning-disk confocal microscope (Figure 4A). Visualization of the inocula showed that approximately 88% of HIV-1_NL4-3/Gag-iVenus_ contained mScarlet-Vpr (Figure S8A). HIV-1_NL4-3/Gag-iVenus_ containing mScarlet-Vpr showed reduced but still substantial infectivity relative to HIV-1_NL4-3_ (Figure S8B). We observed colocalization of ∼55% of HIV-1 particles with BSA-AlexaFluor647 (Figure 4B, white arrowheads). The EIPA treatment prevented the uptake of BSA-AlexaFluor647, resulting in significantly suppressed colocalization of HIV-1 particles with BSA-AlexaFluor647 (Figure 4B and C). HIV-1 internalization via macropinocytosis was insensitive to T-20 (Figure 4C), which is consistent with the notion that fusion-arrested virus particles can be endocytosed (Figure 2). These results indicate that both macropinocytosis and non-macropinocytic endocytosis serve as a pathway for HIV-1 internalization into primary CD4^+^ T cells and suggest that EIPA dampens HIV-1 internalization through macropinocytosis into primary CD4^+^ T cells.

**Figure 4.**
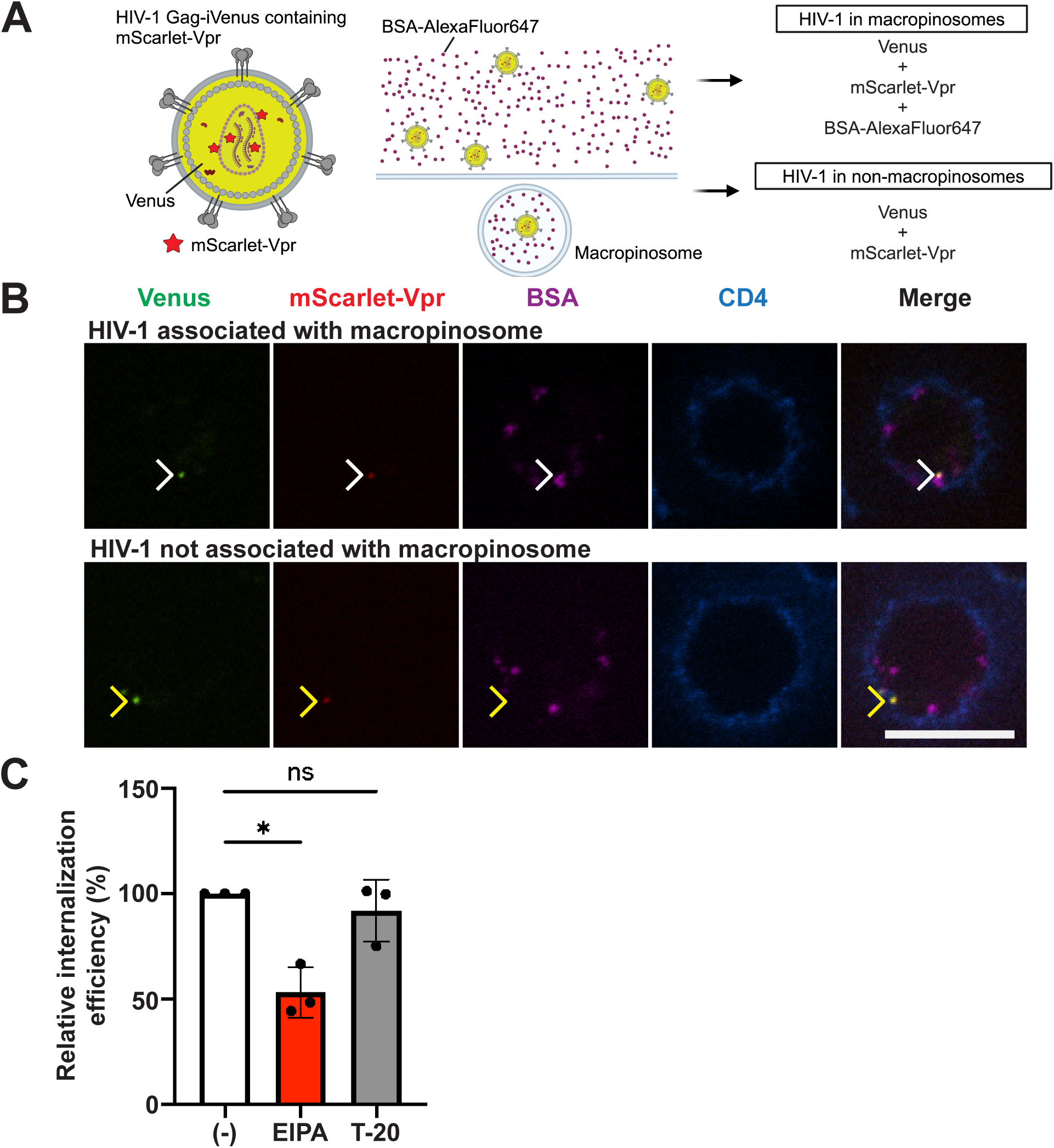
Obsaervation of macropinocytosis-mediated HIV-1 internalization into primary CD4^+^ T cells by a microscopic analysis. (A) A schematic illustration of the experimental procedure performed for panels B and C. Activated primary CD4^+^ T cells were pretreated with either EIPA, T-20, or vehicles for 15 minutes at 37°C and inoculated with HIV-1_NL4-3/Gag-iVenus_ containing mScarlet-Vpr in the presence of BSA-AlexaFluor647 and in the presence or absence of either EIPA or T-20. Inoculated cells were trypsinized, labeled with an anti-CD4 antibody conjugated with BV421, plated on top of a poly-L-lysin-coated glass bottom chamber, and fixed. The fixed cells were observed with a spinning-disk confocal microscope. (B) Examples of internalized Venus- and Vpr-positive HIV-1 particles detected in individual *z*-slices of cells treated with vehicles are shown. White and yellow arrowheads indicate an HIV-1 particle colocalized with BSA-AlexaFluor647 ("HIV-1 in macropinosomes") and an HIV-1 particle not colocalized with BSA-AlexaFluor647 ("HIV-1 in non-macropinosomes"), respectively. Bars, 10 μm. (C) Quantification of the relative internalization efficiency. The punctate signals for internalized HIV-1 in macropinosomes (panel B) were quantified and normalized with cell numbers. A total of more than 51 cells in each condition were examined in each of three independent experiments. The *p* values were determined using Dunnett’s test following one-way ANOVA. *, *p*<0.05; n.s., not significant.

### A microscopy observation showed HIV-1 fusion at macropinosomes in primary CD4^+^ T cells

Even though it has been reported that HIV-1 fusion takes place at endosomal compartments in primary CD4^+^ T cells (12–14), which endocytic pathway(s) mediate HIV-1 fusion is unknown. To investigate whether HIV-1 fusion takes place at macropinosomes, we attempted to use HIV-1_NL4-3_ with membranes labeled by lipophilic dyes, R18 and DiOC18. It has been reported that influenza virus labeled with this combination allows for detection of the subcellular fusion sites based on the increase in emission of the DiOC18 fluorescence due to dye dequenching (37). We observed the co-localization of macropinosomes and labeled HIV-1 in primary CD4^+^ T cells in two independent experiments. However, in the two different conditions that we examined, we observed the dequenching of DiOC18 at macropinosomes even in the presence of a fusion inhibitor, AMD3100 (Fig. S9). Although these results indicate that non-engineered HIV-1 is internalized into primary CD4+ T cells via macropinosomes, they also suggest that this labeling method is not applicable to examine the site of HIV-1 fusion. As an alternative approach, we decided to use HIV-1_NL4-3/MA-3xHA_, which encodes three copies of an HA-tag near the C terminus of the MA domain of Gag (MA-3xHA) (38) (Figure 5A). Since MA should remain at sites of HIV-1 fusion because of its membrane-binding ability (39), the HA signal associated with macropinosomes would represent either the fusion at the macropinosomes or pre-fusion virus particles therein. The insertion of 3xHA within the C-terminal region of MA did not affect virus assembly, release, and maturation (Figure S10). The infectivity of HIV-1_NL4-3/MA-3xHA_ was approximately 60% of that of HIV-1_NL4-3_ (Figure S8B).

**Figure 5.**
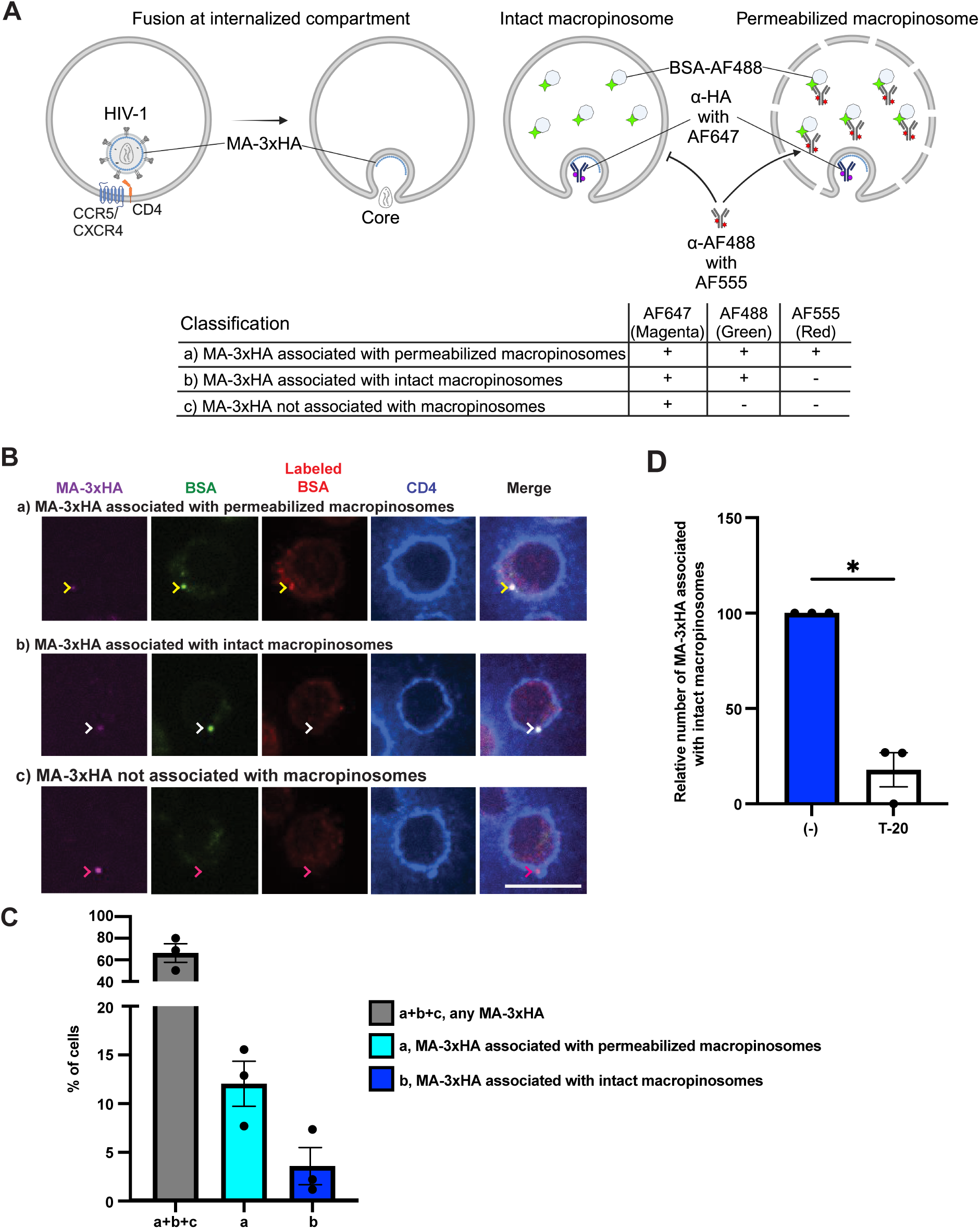
HIV-1 fusion occurs at macropinosomes in primary CD4^+^ T cells. (A) A schematic illustration of the experimental procedure performed for panels B and C. Activated primary CD4^+^ T cells were inoculated with HIV-1_NL4-3/MA-3xHA_ in the presence of BSA-AlexaFluor488. Cells were labeled with an anti-CD4 antibody conjugated with BV421, plated on top of a poly-L-lysin-coated glass bottom chamber, and fixed. The fixed cells were permeabilized with digitonin and labeled with an anti-AlexaFluor488 antibody followed by the secondary antibody conjugated with AlexaFluor555 and an anti-HA antibody conjugated with AlexaFluor647. The cells were observed with a spinning-disk confocal microscope. (B) Individual *z*-slices of representative cells are shown. Yellow arrowheads indicate MA-3xHA signals colocalized with BSA-AlexaFluor488 and AlexaFluor555 (permeabilized macropinosome with MA-3xHA), white arrowheads indicate MA-3xHA signals colocalized with BSA-AlexaFluor488 without AlexaFluor555 (intact macropinosome with MA-3xHA), and pink arrowheads indicate MA-3xHA signals colocalized with neither BSA-AlexaFluor488 nor AlexaFluor555. Bars, 10 μm. (C) Quantification of the percentages of cells with any MA-3xHA, a+b+c; MA-3xHA associated with permeabilized macropinosomes, a; and MA-3xHA associated with intact macropinosomes, b. (D) Quantification of the number of MA-3xHA puncta associated with intact macropinosomes. The number of MA-3xHA puncta was normalized with cell numbers. The experiments were performed three times with primary CD4^+^ T cells isolated from three donors. A total of more than 134 cells were examined in each of three independent experiments. The *p* values were determined using two-tailed paired Student′s *t*-test. *, *p*<0.05.

Activated CD4^+^ T cells were inoculated for 2 hours with HIV-1_NL4-3/MA-3xHA_ in the presence of BSA-AlexaFluor488 and in the presence or absence of T-20. CD4 on the surface of cells was labeled to identify the cell surface. To detect the post-fusion MA-3xHA, which should be exposed at the cytoplasmic side of the macropinosome membrane, while minimizing the detection of pre-fusion viruses inside macropinosomes, we treated the fixed cells with a very low concentration of digitonin to permeabilize only the plasma membrane but not the macropinosome and viral membranes prior to immunostaining with an anti-HA antibody conjugated with AlexaFluor647. Nonetheless, since both macropinosome and viral membranes originate from the plasma membrane, it is possible that digitonin permeabilizes those membranes in addition to the plasma membrane, thereby making MA-3xHA in pre-fusion virus particles within macropinosomes accessible to the anti-HA antibody. To distinguish MA-3xHA present at the cytoplasmic side of macropinosomes (indicating HIV-1 fusion) from MA-3xHA exposed due to permeabilization of both macropinosomes and viral membranes, we used an anti-AlexaFluor488 antibody followed by a secondary antibody conjugated with AlexaFluor555, which will detect BSA-AlexaFluor488 in permeabilized macropinosomes but not intact macropinosomes. Based on distances between the fluorescence signals, which were measured using the Imaris software (Figure S11), we classified MA-3xHA signals into three groups, a) associated with permeabilized macropinosomes, b) associated with intact macropinosomes, and c) not associated with macropinosomes (Figure 5A and B). We quantified the percentages of total cells with MA-3xHA, cells with MA-3xHA associated with permeabilized macropinosomes, and cells with MA-3xHA associated with intact macropinosomes, i.e., HIV-1 fusion at macropinosomes. Among the total cells examined, ∼66% of primary CD4^+^ T cells displayed MA-3xHA signals, which includes cells showing MA-3xHA signals that do not colocalize with either BSA-AlexaFluor488 or AlexaFluor555 (Figure 5B, MA-3xHA not associated with macropinosomes and Figure 5C, a+b+c). In ∼12% of total cells, we observed the association of MA-3xHA with permeabilized macropinosomes (Figure 5B, MA-3xHA associated with permeabilized macropinosomes, and Figure 5C, a), which includes signals representing HIV-1 fusion at macropinosomes and pre-fusion virus particles therein. Finally, approximately 3.5% of total cells showed the association of MA-3xHA with intact macropinosomes (Figure 5B, MA-3xHA associated with intact macropinosomes, and Figure 5C, b). We also quantify the number of MA-3xHA puncta associated with intact macropinosomes. Importantly, we found that the T-20 treatment inhibited the exposure of MA-3xHA at intact macropinosomes (Figure 5D). These results demonstrated that HIV-1 fusion can occur at macropinosomes in primary CD4^+^ T cells.

### Macropinocytosis inhibitors inhibit HIV-1 infection of primary CD4^+^ T cells

Our group previously showed that macropinocytosis is essential for sustaining mTORC1 activity in CD4^+^ T cells(17, 19). Additionally, other groups reported that active mTORC1 regulates HIV-1 infection(40–46). Therefore, it is conceivable that inhibition of macropinocytosis can prevent HIV-1 infection through the inhibition of both entry step and mTORC1 activation. We first sought to determine whether EIPA inhibits mTORC1 activity in primary CD4^+^ T cells. Activated primary CD4^+^ T cells were treated or not with either EIPA, or AZD2014, which is a specific mTOR inhibitor, or T-20 for 2 hours and 15 min at 37°C. The treatment of primary CD4^+^ T cells with either EIPA or AZD2014 inhibited the phosphorylation of the S6 protein, indicating that these compounds suppress mTORC1 activity in primary CD4^+^ T cells (Fig. 6A and B). To investigate the effect of these compounds on productive infection by HIV-1, activated primary CD4^+^ T cells and A3.01 cells were inoculated with a replication-competent HIV-1_NL4-3_ encoding NanoLuc (HIV-1_NL-NI_) for 2 hours at 37°C in the presence or absence of EIPA, AZD2014, or T-20. Cells were then washed to remove unbound viruses and compounds and further cultured for 24 hours at 37°C in the presence of SQV to prevent multiple-cycle replication (Figure 6C). EIPA but not AZD2014 inhibited HIV-1_NL-NI_ infection of primary CD4^+^ T cells (Figure 6D). In contrast, neither EIPA nor AZD2014 inhibited HIV-1_NL-NI_ infection of A3.01 cells (Figure 6E), even though EIPA partially inhibited HIV-1 fusion with this cell type (Figure 3G). In addition, the infection of primary CD4^+^ T cells with a transmitted/founder virus encoding NanoLuc (HIV-1_CH040-NI_) was inhibited by EIPA more efficiently than AZD2014 (Figure 6F). These results indicate that EIPA inhibits HIV-1 infection of primary CD4^+^ T cells independently of the coreceptor usage. Considering that the inhibitory effect of EIPA on HIV-1 infection was slightly stronger than the effect on HIV-1 internalization and fusion (compare Figures 2 and 3 versus 6), these results suggest that in addition to HIV-1 entry, EIPA may inhibit other step(s) of HIV-1 replication cycle in primary CD4^+^ T cells, but it appears unlikely to be a step involving mTORC1 activity. In addition to EIPA, J/B inhibited HIV-1_NL-NI_ infection of primary CD4^+^ T cells (Figure S12). Altogether, these results demonstrated that inhibitors of macropinocytosis prevent HIV-1 infection of primary CD4^+^ T cells.

**Figure 6.**
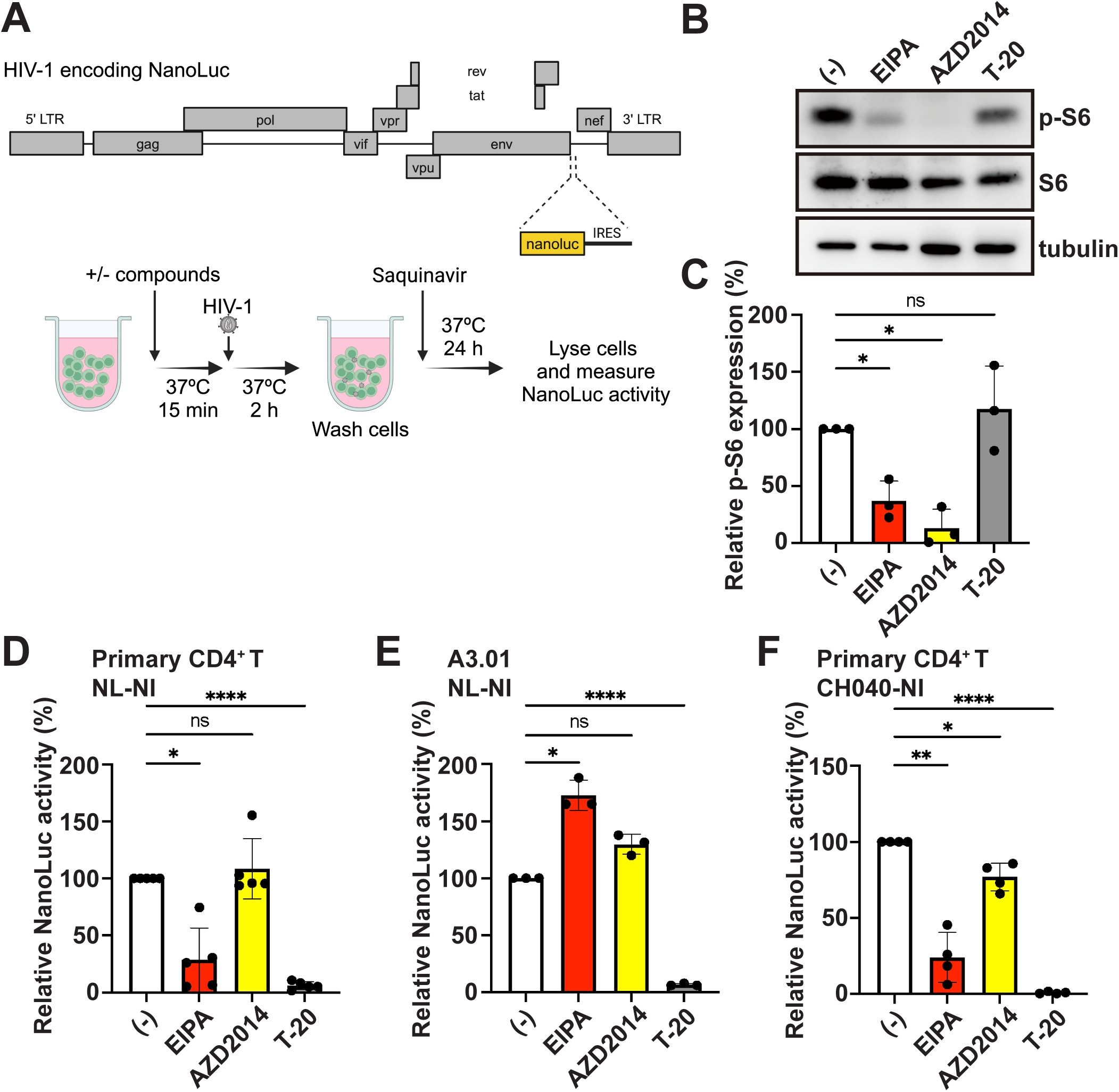
Inhibition of macropinocytosis diminishes HIV-1 infection of primary CD4^+^ T cells. (A) Analyses of mTORC1 activity by detection of the phosphorylated S6 (p-S6) protein. Activated primary CD4^+^ T cells were treated with either EIPA, AZD2014, T-20, or vehicles for 2 hours and 15 min at 37°C. Phosphorylated S6, total S6, and tubulin expression in cell lysates was examined by western blotting. The representative images were shown. (B) Quantification of the relative expression of the p-S6 normalized by the expression of total S6 protein. (C) A schematic illustration of the experimental procedure of D-F. Activated primary CD4^+^ T cells and A3.01 cells were pretreated with either EIPA, AZD2014, T-20, or vehicles for 15 min at 37°C and inoculated with HIV-1 encoding NanoLuc for 2 hours at 37°C. Cells were washed and cultured for 24 hours at 37°C in the presence of saquinavir. Cells were then lysed, and the NanoLuc activity in the cell lysates was measured. The NanoLuc activity was normalized by total protein amounts. (D and F) Indicated cells were inoculated with indicated viral strains. Data are presented as mean +SD in panel B and mean ±SD in panels D-F, respectively. The experiments were performed with primary CD4^+^ T cells isolated from three (A and B) and five (D and F) donors, respectively, in three independent experiments. The experiments were performed three times with A3.01 cells independently in E. The *p* values were determined using Dunnett’s test following one-way ANOVA in B, and D-F. *, *p*<0.05; **, *p*<0.01; ****, *p*<0.0001; n.s., not significant.

## Discussion

While the contribution of different endocytic pathways to productive infection has been shown (5–8, 12–14), the exact subcellular location of HIV-1 entry into primary CD4^+^ T cells, which is the major target of HIV-1, remains to be determined. In the present study, we showed using HIV-1 bearing SOS Env that HIV-1 entry into primary CD4^+^ T cells can take place at both the plasma membrane and internalized compartments. Under the same condition, HIV-1 entered two commonly used T cell lines, A3.01 cells and Jurkat cells, primarily at the plasma membrane. We further observed that EIPA and J/B, macropinocytosis inhibitors, inhibited HIV-1 internalization into, fusion with, and infection of primary CD4^+^ T cells. We also detected HIV-1 fusion at macropinosomes, endosomes derived from macropinocytosis, in primary CD4^+^ T cells. Altogether, these results highlight the important role of macropinocytosis in HIV-1 infection of primary CD4^+^ T cells. This study additionally emphasizes the importance of using primary CD4^+^ T cells, as opposed to T cell lines, as HIV-1 target cells to elucidate the detailed molecular mechanisms of HIV-1 entry.

Primary CD4^+^ T cells engage in multiple endocytic pathways, e.g., clathrin-mediated endocytosis, non-clathrin-mediated endocytosis, and macropinocytosis (16). Using microscopic approaches, we demonstrate that macropinocytosis contributes to internalization into and subsequent fusion of HIV-1 with primary CD4^+^ T cells. This is consistent with the previous observations that are apparently contradictory to each other: one showed that HIV-1 enters into primary CD4^+^ T cells at the plasma membrane but not endosomes formed at 22°C (15) and another that HIV-1 entry occurs at endosomal compartments in primary CD4^+^ T cells (12–14). The observation that macropinocytosis is sensitive to the reduced temperature at which other endocytosis activities can persist (Figure S2) reconciles the hitherto unresolved discrepancy over the role of endocytosis in HIV-1 infection of primary CD4^+^ T cells (see Introduction).

To determine the location of HIV-1 fusion, we used HIV-1 encoding MA-3xHA because MA can remain on the cytoplasmic side of the membrane at the site of fusion (39). In addition, to avoid the detection of pre-fusion viral particles inside macropinosomes as much as possible, we used digitonin to permeabilize the plasma membrane. This experimental approach has several limitations: i) HIV-1 fusion at the plasma membrane is not detectable because upon HIV-1 fusion with the plasma membrane, MA-3xHA is likely to diffuse laterally from the site of fusion on the plasma membrane, resulting in a loss of signals; ii) The identity of the endosomal compartments associated with MA-3xHA signals not associated with macropinosomes and whether these HA signals represent pre- or post-fusion virus cannot be determined; iii) Even though we used very low concentration of digitonin to permeabilize only the plasma membrane, some macropinosomes were permeabilized. Therefore, although MA-3xHA signals associated with intact macropinosomes represent sites of fusion, the fraction of cells with these signals is likely an underestimation of the fusion events at macropinosomes. Additionally, since HIV-1 fusion at the plasma membrane cannot be detected using this approach, the relative fraction of fusions at macropinosomes versus other locations remains to be determined. Despite these limitations, the results obtained with MA detection in digitonin-permeabilized cells strongly support the notion that HIV-1 fusion occurs in macropinosomes in primary CD4^+^ T cells. To determine the fraction of HIV-1 fusion that occurs at macropinosomes, live-cell imaging that allows for tracking of internalized particles up to the point of content release should be applied (5, 12, 13) in conjunction with macropinosome markers, such as BSA.

HIV-1 fusion with primary CD4^+^ T cells was recently reported to take place at endosomes with neutral pH (12). Considering that macropinosomes are acidified during their maturation similar to other endosomes (20, 47), HIV-1 fusion might occur early after macropinocytosis. The acidification of macropinosomes is likely derived from the fusion with lysosomes. In murine bone marrow-derived macrophages, macropinosome-lysosome fusion happens in 7-8 minutes after the formation of the macropinosome (48). This time scale would dovetail with the observation that the half-time for HIV-1 fusion within the neutral-pH endosomes of primary CD4^+^ T cells is approximately 16 minutes (12). Nonetheless, it is possible that macropinocytosis in primary CD4^+^ T cells may differ from that in murine bone marrow-derived macrophages, particularly concerning the timing of acidification. Additionally, there exists the possibility that HIV-1 selectively exploits macropinosomes that evade acidification. Macropinosomes can occasionally recycle back to the plasma membrane without fusing to lysosomes (49), and these macropinosomes might mediate HIV-1 entry into primary CD4^+^ T cells. Elucidating the properties of macropinosomes containing HIV-1 particles, such as intravesicular pH and their intracellular trafficking pathways, in primary CD4^+^ T cells is critical for understanding the mechanisms underlying HIV-1 fusion within neutral pH endosomes.

Our experiments using HIV-1 with SOS Env and the membrane-impermeable fusion inhibitor T-20 suggest that the sites of HIV-1 fusion with primary CD4^+^ T cells are at both the plasma membrane and internal compartments formed at 37°C. One of the limitations of SOS Env is that the infection efficiency of HIV-1 with SOS Env induced by DTT is unlikely comparable with that of HIV-1 with wild-type Env (50). The accurate comparison of infectivity between wild-type Env and DTT-treated SOS Env in cell-based assays is not feasible because SOS Env requires receptor and coreceptor engagement prior to DTT treatment (30). In addition to this limitation, it is possible that SOS Env-bearing HIV-1 on the surface of target cells can be more efficiently internalized into target cells prior to DTT treatment due to the fusion arrest than HIV-1 with wild-type Env. This potential enhancement of SOS Env-containing HIV-1 internalization may bias our analysis towards overestimation of the fraction of cytoplasmic HIV-1 entry through internalized compartments. Notably, however, at least in this approach, we observed the cell type-specific difference in the subcellular locations of HIV-1 entry between primary CD4^+^ T cells and two T cell lines.

There is an apparent discrepancy regarding the results obtained with A3.01 cells. Consistent with the observation obtained with SOS Env (Figure 1), EIPA does not inhibit productive infection of this cell line by HIV-1 and rather enhances infection (Figure 6E). However, the BlaM-Vpr-based fusion assay demonstrates that EIPA attenuates HIV-1 fusion with A3.01 cells (Figure 3G). This supports the possibility that in A3.01 cells, the EIPA-sensitive fraction of HIV-1 fusion, presumably taking place at macropinosomes, does not contribute to productive infection. Considering that EIPA enhances productive infection of A3.01 cells by HIV-1 without apparent effect on HIV-1 internalization into A3.01 cells as measured by the NanoLuc-based assays (Figure 2C), it is conceivable that in EIPA-treated A3.01 cells, productive fusion at the cell surface is more efficient than in untreated A3.01 cells.

While this manuscript was in preparation, a study reported that EIPA modestly (∼40%) inhibits HIV-1 infection of primary CD4^+^ T cells (12). In the current study, we observed a more robust inhibition of both CXCR4- and CCR5-using HIV-1 infection of primary CD4^+^ T cells using a similar EIPA treatment protocol (∼75%). This difference may be explained by a difference in inoculation procedures: simple inoculation in the current study versus spinoculation in the published study. Spinoculation, disrupts the actin cytoskeleton (51). Therefore, spinoculation may have impeded macropinocytosis, as well as other non-macropinocytic endocytosis pathways that are regulated by actin (52). Consequently, the efficiency of HIV-1 macropinocytosis and other actin-dependent endocytic processes involved in HIV uptake may be affected. However, the impact of spinoculation on the endocytotic activities in primary T cells remains to be determined.

Since the inhibition of internalization by EIPA is 30∼50% in our experimental setting, the inhibition of macropinocytosis by EIPA might attenuate additional steps of the HIV-1 replication cycle in primary CD4^+^ T cells. One possible mechanism by which EIPA inhibits HIV-1 infection is suppression of mTORC activity. mTORC activity correlates with efficient HIV-1 infection of primary CD4^+^ T cells (43–46). However, AZD2014, a specific mTOR inhibitor that inhibits mTORC1 and mTORC2 activities, shows no or only a modest inhibitory effect on HIV-1 infection compared to EIPA in our experimental setting. Therefore, regardless of the involvement of mTORC, the primary effect of EIPA on HIV-1 infection is likely the inhibition of the cytoplasmic entry process of HIV-1, and this effect of EIPA is evident only in primary CD4^+^ T cells but not in A3.01 cells.

HIV-1 uses macropinocytosis to enter primary CD4^+^ T cells (shown by this study) and other cells (21–24). In addition to HIV-1, multiple virus species enter target cells via macropinocytosis (25–29). These observations suggest a possibility that macropinosomes provide beneficial environments for some viruses to enter target cells compared to the plasma membrane and other endosomal compartments. For example, pro-viral factors, e.g., viral receptors, may accumulate on macropinosome membranes. Macropinosomes contain HIV-1 coreceptor CCR5 and likely CXCR4 (53, 54). In addition, HIV-1 fusion within macropinosomes would require the presence of CD4 on the lumen surface of these structures, consistent with the observation that CD4 is present in forming macropinosomes (19). CD4, CCR5, and CXCR4 may be highly concentrated on the macropinosome membranes in primary CD4^+^ T cells compared to the plasma membrane and/or other endosomal compartments. Inversely, it is also possible that antiviral factors, such as interferon-inducible transmembrane (IFITM) proteins, which localize to endosomes (55–57), may be excluded from macropinosomes. Specific membrane proteins are observed to be excluded from macropinosomes in *Dictyostelium* amoebae (58). This indicates the presence of a mechanism for the exclusion of specific proteins from macropinosomes during macropinocytosis. Whether proviral and antiviral host membrane proteins are enriched or excluded from macropinosomes remains to be determined.

In summary, we uncovered that macropinocytosis contributes to HIV-1 entry into primary CD4^+^ T cells, which is not evident in T cell lines. Since macropinocytosis in primary CD4^+^ T cells was only recently discovered, the molecular mechanisms are not well characterized. While the molecular mechanisms of macropinocytosis are similar across different cell types examined thus far, there are also significant differences (16–19). Fully understanding the molecular mechanism promoting macropinocytosis in primary CD4^+^ T cells may identify a novel target for an antiretroviral strategy that specifically targets macropinocytosis in this cell type without adverse effects on other cell types.

## Methods

### Human cell lines

Human T cell lines, A3.01 cells and Jurkat cells, obtained from NIH AIDS Reagent Program were cultured in Roswell Park Memorial Institute (RPMI) medium (Invitrogen) supplemented with 10% fetal bovine serum (FBS, Invitrogen) and 50 units/mL penicillin-streptomycin (Gibco) at 37°C. Human 293T cells obtained from ATCC and TZM-bl cells obtained from NIH AIDS Reagent Program were grown in Dulbecco’s modified eagle medium (DMEM, either Lonza or Corning) supplemented with 10% FBS and 50 units/mL penicillin-streptomycin at 37°C.

### Human primary cells

Peripheral blood mononuclear cells (PBMCs) were obtained from buffy coats derived from healthy donors (New York Blood Center) and cryopreserved when PBMCs were not used for further experiments immediately. Primary CD4^+^ T cells were isolated from PBMCs by the negative selection using the CD4^+^ T cell isolation kit (Miltenyi Biotec). To activate PBMCs with PHA-P (Sigma), PBMCs were cultured in RPMI containing 10% FBS, 1% penicillin-streptomycin, 6 μg/mL PHA-P, and 20 units/mL human recombinant IL-2 (Roche) for 44 hours at 37°C. To activate primary CD4^+^ T cells and PBMCs with anti-CD3/CD28 antibodies, 0.5 μg/mL of an anti-CD3 antibody (clone; OKT3, eBioscience) in 1xPBS(-) was added to the 6-well culture plate and incubated for at least 2 hours at 37°C. The plate was washed with 1xPBS(-) twice to remove unbound antibodies. Purified CD4^+^ T cells or PBMCs were resuspended with RPMI medium supplemented with 10% FBS, 1% penicillin-streptomycin, 20 units/mL human recombinant IL-2, and 1 μg/mL an anti-CD28 antibody (clone; 28.2, eBioscience). Cells were then plated onto the anti-CD3 antibody-coated plate and cultured for 44 or 68 hours at 37°C for activation.

### Plasmids

A lab-adapted HIV-1 molecular clone pNL4-3 and a transmitted/founder HIV-1 molecular clone, pCH040 (59), were obtained from NIH AIDS Reagent Program. A replication-competent HIV-1 molecular clone pNL4-3/Gag-iYFP (Gag-iVenus) was constructed in the previous study (60). A replication-competent HIV-1 molecular clone encoding NanoLuc, pNL-NI, was generated by the restriction enzyme digestion of pCMV6-AC/partial NLenv-NanoLuc-IRES-nef and pNL4-3 at BamHI and XhoI sites and the ligation of these digested plasmids. pCMV6-AC/partial NLenv-nef was generated by the restriction enzyme digestion of pCMV6-AC (Origene) and pNL4-3 at BamHI and XhoI sites and the ligation of these digested plasmids. pCMV6-AC/partial NLenv-NanoLuc-IRES-nef was generated by introducing *nanoluc* and the IRES sequence between *env* and *nef* ORFs by using Gibson Assembly Kit (NEB). The PCR product containing *nanoluc* sequence was obtained with pBD-PASTN (61) as a template. A replication competent transmitted/founder HIV-1 clone, pCH040-NI was generated by introducing *nanoluc* and the IRES sequence between *env* and *nef* ORFs. The PCR fragments containing restriction sites and *nanoluc* and the IRES sequence were prepared by using overlap PCR and digested by StuI/BlpI.

StuI/BlpI-digested PCR fragments were then ligated with StuI/BlpI-digested pCH040. Fusion defective HIV-1 molecular clones, pNL-NI/SOS Env, was constructed by introducing two amino acid substitutions in gp120 and gp41 subunits of Env (A499C and T603C). A part of NL4-3 Env containing the substitutions was amplified by overlap PCR. The PCR product and pNL4-3 were digested by NheI and AleI and ligated to construct pNL4-3/SOS Env. Then, pNL4-3/SOS Env and pNL-NI were digested by SalI and BamHI and ligated to construct pNL-NI/SOS Env. A replication competent HIV-1 molecular clone pNL4-3/Gag-iNanoLuc was constructed by swapping the YFP ORF of pNL4-3/Gag-iVenus with the *nanoluc* ORF. The PCR product containing the *nanoluc* ORF and restriction digestion sequences were amplified by the overlap PCR. The PCR product and pNL4-3/Gag-iVenus were digested by BssHII and XbaI and ligated. A replication competent HIV-1 molecular clone pNL4-3/Gag-iNanoLuc/CH040env was constructed by restriction digest and ligation. pNL4-3/Gag-iNanoLuc and pCH040 were digested by EcoRI and BlpI and ligated. A replication competent HIV-1 molecular clone pNL4-3/MA-3xHA was constructed according to the previous paper (38). An expression plasmid for BlaM-Vpr, pCMV4-BlaM-Vpr (36), was obtained from Addgene. An expression plasmid for mScarlet-Vpr, pmScarlet-Vpr, was generated by the restriction enzyme digestion of pEGFP-Vpr (62) (NIH AIDS Reagent Program) and pmScarlet_C1 (63) (Addgene) with BsrGI and BamHI and the ligation of digested plasmids.

### Virus production

To produce HIV-1_NL4-3/Gag-iNanoLuc_, HIV-1_NL4-3/Gag-iNanoLuc/CH040env_, HIV-1_NL-NI_, HIV-1_CH040-NI_, HIV-1_NL-NI/SOS Env_, and HIV-1_CH040-NI/SOS Env_, 5.25×10^6^ 293T cells were plated on 10 cm tissue culture dishes and cultured for 1 day at 37°C. Cells were then transfected with 20 μg pNL4-3/Gag-iNanoLuc, pNL4-3/Gag-iNanoLuc/CH040env, pNL-NI, pCH040-NI, and pNL-NI/SOS Env using Lipofectamine2000 (Invitrogen). To generate HIV-1_NL4-3_ and HIV-1_CH040_containing BlaM-Vpr, 5.25×10^6^ 293T cells were plated on 10 cm tissue culture dishes and cultured for 1 day at 37°C. Cells were then transfected with 15 μg of either pNL4-3 or pCH040, 5 μg of pCMV4-BlaM-Vpr, and 2.5 μg of pAdvantage (Promega) using Lipofectamine2000. To generate HIV-1_NL4-3/Gag-iVenus_ containing mScarlet-Vpr, 5.25×10^6^ 293T cells were plated on 10 cm tissue culture dishes or 1.05×10^6^ 293T cells were plated on a 6-well culture plate and cultured for 1 day at 37°C. Cells cultured on the 10 cm tissue culture dishes or the 6-well culture plate were then transfected with 15 or 3 μg of either pNL4-3/Gag-iVenus and 5 or 1 μg of pmScarlet-Vpr using Lipofectamine2000. To generate HIV-1_NL4-3_ and HIV-1_NL4-3/MA-3xHA_, 5.25×10^6^ 293T cells were plated on 10 cm tissue culture dishes or 1.05×10^6^ 293T cells were plated on a 6-well culture plate and cultured for 1 day at 37°C. Cells cultured on the 10 cm tissue culture dishes or the 6-well culture plate were then transfected with 20 or 4 μg of either pNL4-3 or pNL4-3/MA-3xHA using Lipofectamine2000. Two days post-transfection, virus-containing supernatants were filtered through a 0.45 µm filter and concentrated by ultracentrifugation at 24,000 rpm for 2 hours at 4°C using a 20% sucrose cushion in a TH641 rotor (ThermoFisher) or SW40Ti (Beckman Coulter). Virions were resuspended with RPMI containing 10% FBS, aliquoted, and stored at −80°C. To quantify virus amounts, the CAp24 amount was measured by enzyme-linked immunosorbent assay (ELISA) (64).

### Labeling HIV-1 particles with R18 and DiOC18

HIV-1_NL4-3_ were labeled with 3,3′-dioctadecyloxacarbocyanine (DiOC18) and octadecyl rhodamine B (R18). Ultracentrifuged HIV-1 was suspended in 5 ml of phosphate-buffered saline. DiOC18 and R18 (Invitrogen) were dissolved in ethanol and mixed at concentrations of 33 μM and 67 μM, respectively. The probe mixture (6 μl) was added to the virus suspension and mixed. The reaction mixture was incubated for 1 hour at room temperature and then used as inoculum.

### ELISA for CAp24

ELISA for CAp24 was performed as described previously (64). In brief, the supernatant containing viral particles was lysed in ELISA lysis buffer [0.05% Tween 20, 0.5% Triton X-100, and 0.5% casein in 1xPBS(-)]. An anti-HIV-1 p24 antibody (clone 183-H12-5C; NIH AIDS Reagent Program) was bound to Nunc MaxiSorp plates overnight at 4°C. Lysed samples were added to the plates and incubated for 2 hours, and captured p24 was detected by sequential incubation with a biotinylated anti-HIV-1 p24 antibody (clone 31–90-25; American Type Culture Collection or clone 241-D; NIH AIDS Reagent Program), streptavidin-HRP (Fitzgerald), and 3,3′,5,5′-tetramethylbenzidine substrate (Sigma). Biotinylation of 31–90-25 and 241-D was performed using the EZ-Link Micro Sulfo-NHS-Biotinylation Kit (Pierce). CAp24 concentrations were determined using the recombinant HIV-1 IIIB p24 protein for standards (NIH AIDS Reagent Program).

### HIV-1 internalization assay

To examine HIV-1 internalization into activated primary CD4^+^ T cells, HIV-1_NL4-3/Gag-iNanoLuc_ and HIV-1_NL4-3/Gag-iNanoLuc/CH040env_ were used. 1×10^6^ activated primary CD4^+^ T cells and A3.01 cells were plated in 96-well U bottom plates. Cells were pretreated with either 50 μM EIPA (Sigma), 1_μM jasplakinolide (Tocris) and 75_μM (S)-(-)-blebbistatin (Tocris), 2 μM T-20 (NIH AIDS Reagent Program), or vehicles for 15 minutes at 37°C and inoculated with HIV-1 (4 μg of p24) for 2 hours at 37°C. Cells were then washed to remove unbound viruses and treated with 0.25% Trypsin (Invitrogen) or 2 mg/mL pronase (Roche) for 1 hour at 4°C to remove surface-bound viruses. RPMI-10 was added to each well to stop the activity of trypsin. Cells were then washed and lysed with 1x Passive Lysis Buffer (PLB; Promega). The NanoLuc activity in the lysates was determined by Nano-Glo® Luciferase Assay System (Promega). The NanoLuc activity was normalized by total protein amounts determined using the Pierce^TM^ 660 nm Protein Assay Reagent (Pierce). To investigate the efficiency of trypsinization at 4°C, 1×10^6^ activated primary CD4^+^ T cells were plated in 96-well U bottom plates, incubated for 15 minutes at 4°C, and inoculated with HIV-1 (4 μg of p24) for 2 hours at 4°C. Cells were then washed to remove unbound viruses and treated or not treated with 0.25% Trypsin (Invitrogen) for 1 hour at 4°C. Cell lysates were prepared, and the NanoLuc activity in the lysates was determined by Nano-Glo® Luciferase Assay System.

### Microscopy-based internalization assay

To observe HIV-1 internalization into activated primary CD4^+^ T cells by the microscopic technique, 1×10^6^ cells were pretreated with either 50 µM EIPA, 2 μM T-20, or vehicles for 15 minutes at 37°C and inoculated for 1 hour with HIV-1_NL4-3/Gag-iVenus_ containing mScarlet-Vpr (4 μg of p24) at 37°C in the presence of a cargo of macropinocytosis, 0.4 mg/mL BSA-AlexaFluor647 (Invitrogen). Subsequently, cells were washed with 1xPBS(-) once, trypsinized with 0.25% Trypsin for 1 hour at 4°C, labeled with an anti-CD4 antibody conjugated with BV421 (1:50 clone; OKT4, BioLegend) in 1xPBS(-) containing 1% BSA (Sigma, FACS buffer) for 30 minutes at 4°C, washed with FACS buffer once, and plated onto the poly-L-lysin-coated 8-well glass-bottom chamber slide for 40 minutes at 4°C. Cells were then fixed with 4% PFA/1xPBS(-) for 30 minutes at room temperature and observed with a spinning-disk confocal microscope (Nikon CSU-X1) in Fluoromount G. Z-series of images were acquired with 150 nm intervals between focal planes. To remove the noise and subtract the background, we used Denoise.ai and the rolling ball correction of Nikon Elements software. By maximum intensity projection, the z-series images composed of 89 focal planes were converted to single plane images. The thresholds of each fluorophore were determined using Huang for mScarlet-Vpr and BSA-AlexaFluor647 and Yen for Venus. These methods are available within the Auto Threshold plugin for the ImageJ software (NIH; downloaded from https://imagej.net/). By using GA3 pipelines of Nikon Element software, we quantified the colocalization of mScarlet-Vpr and Venus first because the pipeline cannot analyze three fluorophores simultaneously. We subsequently analyzed the colocalization between Venus colocalized with mScarlet-Vpr and BSA-AlexaFluor647. The colocalization between mScarlet-Vpr and Venus without overlapping BSA-AlexaFluor647 indicates HIV-1 internalization via non-macropinocytic endocytosis. On the other hand, when three fluorophores overlap, we determined them as HIV-1 particles inside macropinosomes. HIV-1 particles inside macropinosomes and HIV-1 particles not associated with macropinosomes were counted and normalized with total cell numbers. CD4 staining was used to determine the cell number.

### BlaM-Vpr HIV-1 fusion assay and coreceptor density analysis

HIV-1 fusion was analyzed using a previously described BlaM-Vpr-based fusion assay(36). 5×10^5^ activated primary CD4^+^ T cells and A3.01 cells were plated in 96-well U bottom plates. Cells were pretreated with either 50 μM EIPA, 1_μM jasplakinolide (Tocris) and 75_μM (S)-(-)-blebbistatin (Tocris), 2 μM T-20, or vehicles for 15 minutes at 37°C and inoculated with HIV-1_NL4-3_ containing BlaM-Vpr (4 μg of p24) or HIV-1_CH040_ containing BlaM-Vpr (6 μg of p24) for 2 hours at 37°C. Cells were washed with CO_2_-independent media (Gibco) and incubated in CO_2_-independent media containing 1 μM CCF2-AM (Invitrogen) for 1 hour at 15°C. Cells were washed with CO_2_-independent media and CO_2_-independent media containing 2.5 mM probenecid (Sigma) and 10% FBS (development media) subsequently. Cells in development media were incubated overnight at room temperature at dark, washed with FACS buffer, labeled with an anti-CXCR4-APC/Cy7 (1:100, clone 12G5, BioLegend) or CCR5-APC/Cy7 (1:100, clone J418F1, BioLegend) antibody in FACS buffer, fixed, and analyzed by flow cytometry, using an LSR-II Fortessa (Becton Dickinson). Data were analyzed using FlowJo software (TreeStar). To determine the density of CXCR4 and CCR5, the mean fluorescent intensity of APC/Cy7 in CXCR4^+^ or CCR5^+^ cells was measured and divided by the mean values of FSC-A of those cells.

### Microscopy-based fusion assay with HIV-1_NL4-3_ labeled with DiOC18 and R18

To determine the site of HIV-1 fusion with activated primary CD4^+^ T cells by the microscopic technique, 1×10^6^ cells were pretreated with 10 μg/mL AMD3100, or the vehicle for 15 minutes at 37°C and inoculated with HIV-1_NL4-3_ labeled with DiOC18 and R18 at 4°C for 1 hour in the presence of a cargo of macropinocytosis, 0.5 µg/mL BSA-AlexaFluor647 (Invitrogen). Subsequently, cells were incubated for either 30 or 60 minutes at 37°C, labeled with an anti-CD4 antibody conjugated with BV421 (1:100, clone Leu-3, BioLegend) in FACS buffer for 30 minutes at 4°C, and plated onto the poly-L-lysin-coated 35 mm MatTek dishes for 40 minutes at 4°C. Cells were then fixed with 4% PFA+0.2% glutaraldehyde/1xPBS(-) for 30 minutes at room temperature and observed with the spinning-disk confocal microscope. Z-series of images were acquired with 200 nm intervals between focal planes. The collected 3D images were analyzed using ImageJ software.

### Microscopy-based fusion assay using HIV-1_NL4-3/MA-3xHA_

To determine the site of HIV-1 fusion with activated primary CD4^+^ T cells by the microscopic technique, 1×10^6^ cells were pretreated with 2 μM T-20 or the vehicle for 15 minutes at 37°C and inoculated with HIV-1_NL4-3/MA-3xHA_ (4 μg of p24) at 37°C for 2 hours in the presence of a cargo of macropinocytosis, 0.4 mg/mL BSA-AlexaFluor488 (Invitrogen). Subsequently, cells were washed with 1xPBS(-) once, labeled with an anti-CD4 antibody conjugated with BV421 (1:100, clone Leu-3, BioLegend) in FACS buffer for 30 minutes at 4°C, washed with FACS buffer once, and plated onto the poly-L-lysin-coated 8-well glass-bottom chamber slide for 40 minutes at 4°C. Cells were then fixed with 4% PFA/1xPBS(-) for 30 minutes at room temperature, permeabilized with 2 μg/mL digitonin in 1xPBS(-) for 1 minute at room temperature, and labeled with an anti-AlexaFluor488 antibody (1:50, Invitrogen A-11094) followed by an anti-rabbit IgG-AlexaFluor555 PLUS (1:1,000, Invitrogen) and an anti-HA-AlexaFluor647 antibody (1:200, clone 6E2, Cell Signaling), and observed with the spinning-disk confocal microscope in Fluoromount G. Z-series of images were acquired with 100 nm intervals between focal planes. A total of 130 focal planes were imaged. The collected 3D images were processed using Imaris software. In this process, fluorescent signals for AlexaFluor488, AlexaFluor555, and AlexaFluor647 were segmented into identifiable spots. We analyzed the percentages of MA-3xHA associated with either permeabilized or intact macropinosomes by using the Shortest Distance algorithm available in Imaris. We measured the radii of the BSA-AlexaFluor488 spots and identified the radius (R_488_) that equals or exceeds radii of 95% of BSA-AlexaFluor488 spots. Subsequently, we calculated the shortest centroid-to-centroid distances from each BSA-AlexaFluor488 spot to the nearest AlexaFluor555 spot (SD_488-555_) and from each BSA-AlexaFluor488 spot to MA-3xHA spot (represented by AlexaFluor647) (SD_488-647_).

BSA-AlexaFluor488 spots with SD_488-555_> R_488_ and SD_488-647_≦ R_488_ were classified as potential intact macropinosomes associated with MA3xHA. On the other hand, BSA-AlexaFluor488 spots with SD_488-555_ ≦ R_488_ and SD_488-647_ ≦ R_488_ were categorized as potential permeabilized macropinosomes associated with MA-3xHA. We then determine whether each potential intact or permeabilized macropinosome is truly intact or permeabilized and whether they are associated with MA-3xHA, both based on the radius of each BSA-AlexaFluor488 spot, SD_488-555_, and SD_488-647_. If the radius is shorter than SD_488-555_ and longer than SD_488-647_, the macropinosome is intact and associated with MA-3xHA. We calculated the percentage of MA-3xHA^+^ primary CD4^+^ T cells and primary CD4^+^ T cells displaying either intact or permeabilized macropinosomes associated with MA-3xHA. Additionally, we counted the number of intact macropinosomes associated with MA-3xHA and normalized the number with cell numbers to examine the effect of T-20. CD4 staining was used to determine the cell number.

### Single-cycle HIV-1 infection assay

For analysis of infection of HIV-1 SOS Env, 1×10^6^ activated primary CD4^+^ T cells, A3.01 cells, and Jurkat cells were plated in 96-well U bottom plates. Cells were inoculated with HIV-1 SOS Env (4 μg of p24) for 2 hours at 37°C. The inoculum was replaced with fresh RPMI containing 10% FBS, 1% penicillin-streptomycin, and 10 mM DTT (Sigma). Cells were incubated for 10 minutes at room temperature (22∼23°C) and washed with 1xPBS(-) three times to remove DTT. Cells were then incubated for 15 minutes at room temperature (22∼23°C) and further cultured at 37°C in the presence of 10 μM SQV (NIH AIDS Reagent Program) and either 15 μM Ral (NIH AIDS Research and Reference Reagent Program) or 2 μM T-20. SQV was added to prevent multiple cycle replication. After 3 days, cells were washed with 1xPBS(-) and lysed with 1xPLB. The NanoLuc activity was analyzed as described above. The NanoLuc activity was normalized by total protein amounts determined with the Pierce^TM^ 660 nm Protein Assay Reagent.

For analysis of infection by replication competent viruses, 1×10^6^ primary CD4^+^ T cells and A3.01 cells were plated in 96-well U bottom plates. Cells were pretreated with either 50 μM EIPA, 1_μM jasplakinolide (Tocris) and 75_μM (S)-(-)-blebbistatin (Tocris), 5 μM AZD2014 (Selleckchem), 2 μM T-20, or vehicles for 15 minutes at 37°C and inoculated with HIV-1 (4 μg of p24) for 2 hours at 37°C. Cells were then washed to remove unbound viruses and compounds three times with 1xPBS(-) and cultured at 37°C in the presence of 10 μM SQV to prevent multiple cycle replication. At 24 hours postinfection, cells were washed with 1xPBS(-) and lysed with 1xPLB. The NanoLuc activity was analyzed as described above.

### Macropinocytosis efficiency analysis

To determine the macropinocytosis efficiency at 22°C, 1×10^6^ activated CD4^+^ T cells were plated in 96-well U bottom plates and incubated for 2 hours at either 4, 22, or 37°C in the presence of 4 μg/mL BSA-AlexaFluor680 (Invitrogen), a cargo of macropinocytosis. Cells were washed, fixed with 4% PFA/1xPBS(-) for 30 minutes at room temperature, and analyzed for BSA uptake by flow cytometry using an LSR-II Fortessa. To investigate the effect of EIPA on macropinocytosis efficiency, 1×10^6^ activated CD4^+^ T cells and A3.01 cells were pretreated with EIPA or the vehicle for 15 minutes at 37°C. 0.4 mg/mL BSA-AlexaFluor680 was added to the cells and incubated for 2 hours at 37°C. As a control, cells were incubated for 15 minutes at either 4 or 37°C in the absence of EIPA and incubated for 2 hours at either 4 or 37°C in the presence of BSA-AlexaFluor680. Cells were washed and analyzed for BSA uptake by flow cytometry using an LSR-II Fortessa. Data were analyzed using FlowJo software.

### mTORC1 activity analysis

One million activated CD4^+^ T cells were plated in 96-well U bottom plates and treated with either 50 μM EIPA, 5 μM AZD2014, 2 μM T-20, or vehicles for 2 hours and 15 minutes at 37°C. Cells were washed and lysed. The lysates were subjected to sodium dodecyl sulfate polyacrylamide gel electrophoresis (SDS-PAGE), followed by immunoblotting using anti-phosphorylated S6 (1:1,000; clone D57.2.2E, Cell Signaling), S6 (1:1,000; clone 5G10, Cell Signaling), and tubulin (1:4,000; clone B-5-1-2, Sigma) antibodies followed by corresponding secondary antibodies conjugated to horseradish peroxidase: anti-mouse IgG (1:20,000; Millipore) and anti-rabbit IgG (1:100,000; Invitrogen) antibodies. Signals were detected using SuperSignal West Pico PLUS or West Atto chemiluminescent substrates (Pierce). To quantify the relative expression of phosphorylated S6, the expression of phosphorylated S6 was normalized by the expression of total S6 protein.

### Toxicity assay

The toxicity of EIPA to primary CD4^+^ T cells was determined by using the APC Annexin V Apoptosis Detection Kit with PI (BioLegend) according to the manufacturer’s protocol. 1×10^6^ activated CD4^+^ T cells were plated in 96-well U bottom plates and pretreated with 50-100 μM EIPA for 2 hours and 15 minutes at 37°C. Cells were washed to remove EIPA with 1xPBS(-) three times and cultured in a growth medium for 24 hours at 37°C. Cells were then washed with Cell Staining Buffer (BioLegend) twice and stained with APC Annexin V and PI for 15 minutes at room temperature. After the staining, cells were analyzed for Annexin V binding and PI staining by flow cytometry using FACSCanto (Becton Dickinson). Data were analyzed using FlowJo software.

### CD4 density analysis

8×10^5^ activated PBMCs were plated in 96-well U bottom plates and treated with either EIPA, J/B, or T-20 for 2 hours at 37°C. Cells were washed with 1xPBS(-) once and labeled with anti-CD3-APC/Cy7 or CD3-AlexaFluor647 (1:100; clone UCHT1, BioLegend) and CD4-AlexaFlor647 or CD4-FITC (1:100; clone OKT4, BioLegend) antibodies in FACS buffer for 30 minutes at 4°C. After the labeling, cells were washed, fixed, and analyzed by flow cytometry using FACSCanto. Data were analyzed using FlowJo software. The density of CD4 was measured by the calculation described above.

### CXCR4 density analysis

8×10^5^ activated PBMCs were plated in 96-well U bottom plates and treated with J/B for 2 hours at 37°C. Cells were washed with 1xPBS(-) once and labeled with anti-CD3-AlexaFluor647 (1:100; clone UCHT1, BioLegend), CD4-FITC (1:100; clone OKT4, BioLegend), and CXCR4-PE (1:100; clone 12G5, BioLegend) antibodies in FACS buffer for 30 minutes at 4°C. After the labeling, cells were washed, fixed, and analyzed by flow cytometry using FACSCanto. Data were analyzed using FlowJo software. The density of CXCR4 was measured by the calculation described above.

### Virus release and maturation assay

For analysis of the effect of 3xHA insertion into the stalk region of MA on the virus release and maturation, 4×10^5^ 293T cells were plated on a 12-well tissue culture plate and cultured for 1 day at 37°C. Cells were transfected with either pNL4-3 or pNL4-3/MA-3xHA using Lipofectamine2000 and cultured for 2 days. Cell and virus lysates were prepared as previously described (65). These lysates were subjected to SDS-PAGE, followed by immunoblotting using HIV-Ig (1:1,000; NIH AIDS Reagent Program) or an anti-HA antibody (1:4000; Abcam, Cat# ab9110) followed by corresponding secondary antibodies conjugated to horseradish peroxidase; anti-mouse IgG (1:20,000; Millipore) and anti-rabbit IgG (1:100,000; Invitrogen) antibodies. Signals were detected using SuperSignal West Pico PLUS chemiluminescent substrate (Pierce).

### TZM-bl assay

To measure virion infectivity, 1×10^4^ TZM-bl cells were inoculated with viruses (50 ng of p24 determined by sandwich ELISA using mAb 31–90-25 as detection antibody) and cultured in the presence of 10 μM saquinavir (NIH AIDS Research Reagent Program). At 48 h postinfection, cells were lysed, and the cell lysates were analyzed for luciferase activity using a commercially available kit (Promega).

### Microscopic analysis of virion incorporation of mScarlet-Vpr

HIV-1_NL4-3/Gag-iVenus_ containing mScarlet-Vpr and HIV-1_NL4-3/Gag-iVenus_ produced from 293T cells were plated onto the 35 mm poly-L-lysin-coated MatTek glass bottom culture dishes (MatTek Life Sciences) for 40 minutes at 4°C. Viruses were then fixed with 4% PFA/1xPBS(-) for 30 minutes at room temperature and observed with the spinning-disk confocal microscope. The colocalization efficiency between Venus and mScarlet-Vpr was determined using the Analyze Particles tool in ImageJ. First, all Gag-iVenus fluorescent spots were detected and segmented. A size threshold of 0.05 µm² to infinity was applied to define the minimum spot size. The Regions of Interest (ROIs) corresponding to Gag-iVenus spots from Analyze Particles were then overlaid onto the mScarlet-Vpr spots. The total number of mScarlet-Vpr spots and the mScarlet-Vpr spots colocalized with Gag-iVenus spots were quantified, and the percentage of colocalization was calculated.

### Quantification and statistical analysis

Data and statistical analysis were performed using Prism 10 (GraphPad Software Inc., USA). For two-group comparisons, two-tailed paired Student′s *t*-test was used. Dunnett’s or Bonferroni’s multiple comparisons test following one-way ANOVA was used for multigroup comparisons. The applied statistical analyses, replicates, and exact values of n are reported in the figure legends. Data are presented as mean_±_SD or SEM and were considered statistically significant when the *p-*value was <0.05.

## Data availability

The data that support the findings of this study are available from the corresponding author upon reasonable request.

## Acknowledgments

We thank the members of the Ono laboratory for helpful discussions and critical reviews of the manuscript. We also thank Drs. Kathleen L. Collins and Andrew Mehle for the anti-HIV-1 p24 antibody (clone 31-90-25) and pBD-PASTN, respectively. The following reagents were obtained through the AIDS Research and Reference Reagent Program, Division of AIDS, NIAID, NIH: Panel of Full-Length Transmitted/Founder (T/F) Human Immunodeficiency Virus Type 1 (HIV-1) Infectious Molecular Clones including CH040, pEGFP-Vpr, T-20, raltegravir, saquinavir, HIV-1 IIIB p24 recombinant protein, HIV-1 p24 monoclonal antibody (183-H12-5C), HIV-1 p24 monoclonal antibody (241-D), and TZM-bl cells from Drs. John C. Kappes, Warner C. Greene, Bruce Chesebro, Kathy Wehrly, Susan Zolla-Pazner, and Xiaoyun Wu, and Tranzyme Inc. We thank Drs. Warner C. Greene and Dorus Gadella for pCMV4-BlaM-Vpr and pmScarlet_C1, respectively. This work is supported by NIH grants R01 AI165381 (to A.O. and P.D.K.).

## Author Contributions

T.M., R.d.S.C., K.C., and Y.T. performed the experiments; P.M. and, Y.T.C. provided reagents; E.R. and R.d.S.C. helped with analyses of imaging data; T.M. and A.O. wrote the manuscript with input from J.A.S. and P.D.K; J.A.S., P.D.K., and A.O. were responsible for research supervision, coordination, and strategy.

## Declaration of Interests

The authors declare no competing interests.

---

**Figure S1.** HIV-1 entry into primary CD4^+^ T cells takes place at both the plasma membrane and internalized compartments. (A) The effect of DTT treatment on the total protein amounts in activated primnary CD4^+^ T cells. Data are presented as means ±SD. The experiments were performed seven times with primary CD4^+^ T cells isolated from seven donors. The *p* values were determined using Tukey’s test following one-way ANOVA.***, *p*<0.001; n.s., not significant. (B) The donor-specific differences between the different treatments are clarified for the results presented in Figure 1B.

**Figure S2.** Macropinocytosis of primary CD4^+^ T cells is suppressed at 22°C but not at 37°C. (A) Gating strategy to analyze macropinocytosis efficiency at 4, 22, or 37°C. Activated primary CD4^+^ T cells were pretreated with either EIPA or T-20 for 15 minutes at 37°C. BSA-AlexaFluor680 was added to the cells, and the cells were further incubated for 2 hours at 37°C. After fixation, cells were analyzed for BSA uptake by flow cytometry. One of the representative data is shown. (B). Quantification of the relative BSA uptake in primary CD4^+^ T cells. Data are presented as mean ±SD. The experiments were performed three times with primary CD4^+^ T cells isolated from three donors. The *p* values were determined using Dunnett’s test following one-way ANOVA. *, *p*<0.05; **, *p*<0.01.

**Figure S3.** EIPA suppresses macropinocytosis in primary CD4^+^ T cells and A3.01 cells. (A) Analyses of toxicity of EIPA to primary CD4^+^ T cells. Activated primary CD4^+^ T cells were treated with 50-100 μM EIPA for 2 hours and 15 minutes at 37°C. Cells were washed to remove EIPA with 1xPBS(-) three times and cultured for 24 hours at 37°C. Cells were then stained with APC Annexin V and PI and analyzed for Annexin V binding and PI staining by flow cytometry. (B) Gating strategy to analyze the effect of 50 μM EIPA on macropinocytosis efficiency. Activated primary CD4^+^ T cells were pretreated with either EIPA or vehicle control for 15 minutes at 37°C. BSA-AlexaFluor680 was added to the cells, and the cells were further incubated for 2 hours at 37°C. After fixation, cells were analyzed for BSA uptake by flow cytometry. One of the representative data is shown. (C-D) Quantification of the relative BSA uptake in primary CD4^+^ T cells (C) and A3.01 cells (D). Data are presented as mean ±SD. The experiments were performed three times with primary CD4^+^ T cells isolated from three donors and three times with A3.01 cells. The *p* values were determined using two-tailed paired Student′s *t*-test. *, *p*<0.05.

**Figure S4.** EIPA inhibits internalization of unmodified HIV-1 into primary CD4^+^ T cells. (A) Trypsinization at 4°C removes HIV-1 particles on the surface of primary CD4^+^ T cells. Activated primary CD4^+^ T cells were inoculated with HIV-1_NL4-3/Gag-iNanoLuc_ for 2 hours at 4 °C. After washing, cells were lysed. The NanoLuc activity in cell lysates was measured and normalized by total protein amounts. (B) Activated primary CD4^+^ T cells were pretreated with either EIPA, T-20, or vehicles for 15 minutes at 37°C and inoculated with unmodified HIV-1_NL4-3_ for 2 hours in the presence or absence of either EIPA or T-20. The amounts of the HIV-1 antigen p24 in cell lysates were measured by p24 ELISA. Data are presented as mean ±SD. The experiments were performed three times with primary CD4^+^ T cells isolated from three donors. The *p* values were determined using two-tailed paired Student′s *t*-test. **, *p*<0.01.

**Figure S5.** Both EIPA and J/B inhibits HIV-1 internalization without affecting CD4 expression on primary CD4^+^ T cell surface. (A) Gating strategy to analyze CD4 expression. Activated PBMCs were treated with either EIPA or J/B for 2 hours at 37°C. Cells were labeled with anti-CD3 and CD4 antibodies and fixed. Cells were then analyzed for the expression of surface CD3 and CD4. One of the representative data is shown. (B and C). Quantification of the relative surface CD4 density on primary CD4^+^ T cells treated with either EIPA (B) or J/B (C). The experiments were performed three times with PBMCs isolated from three donors. Data are presented as means ±SD. (D) Activated primary CD4^+^ T cells were pretreated with either J/B or vehicles for 15 minutes at 37°C and inoculated with HIV-1 molecular clones for 2 hours in the presence or absence of either J/B. The NanoLuc activity in cell lysates was measured and normalized by total protein amounts. The experiments were performed with primary CD4^+^ T cells isolated from three donors in three independent experiments. The *p* values were determined using two-tailed paired Student′s *t*-test. **, *p*<0.01; Data are presented as mean ±SD.

**Figure S6.** The reduction of CXCR4 density does not explain the inhibitory effect of a macropinocytosis inhibitor on HIV-1 fusion with primary CD4^+^ T cells. (A-D) Activated primary CD4^+^ T cells and A3.01 cells were pretreated with either EIPA or T-20 for 15 minutes at 37°C and inoculated with HIV-1 containing BlaM-Vpr for 2 hours at 37°C. Inoculated cells were loaded with CCF2-AM at 15°C for 1 hour and incubated overnight at room temperature. The cells were then labeled with either an anti-CXCR4-APC/Cy7 or CCR5-APC/Cy7 antibody and analyzed by flow cytometry. (A and B) Analyses of surface expression of CXCR4 in primary CD4^+^ T cells by flow cytometry. (A) Representative flow plots of surface expression of CXCR4 on primary CD4^+^ T cells were shown. (B) Quantification of the relative density of surface CXCR4 on primary CD4^+^ T cells in the gates shown in (A). (C and D) Analyses of the CCF2 cleavage in cells in the gates shown in (A). (C) Representative flow plots with the percentages of cleaved CCF2^+^ cells are shown. (D) Quantification of the relative percentages of cleaved CCF2^+^ cells (fusion efficiency) in cells in the gates shown in (A). Data are presented as mean ±SD in B and D. The experiments were performed three times with primary CD4^+^ T cells isolated from three donors. The *p* values were determined using Dunnett’s test following one-way ANOVA. **, *p*<0.01; n.s., not significant.

**Figure S7.** J/B inhibits HIV-1 fusion with primary CD4^+^ T cells without affecting CXCR4 density on the cells. (A) Activated PBMCs were treated with J/B or vehicles for 2 hours at 37°C. Cells were labeled with anti-CD3-AlexaFluor647, CD4-FITC, and CXCR4-PE antibodies and fixed. Cells were then analyzed for the expression of surface CXCR4 in CD4^+^ T cells by flow cytometry. Relative surface density of CXCR4 was determined. (B) Activated primary CD4^+^ T cells were pretreated with either J/B or vehicles for 15 minutes at 37°C and inoculated with HIV-1_NL4-3_ containing BlaM-Vpr for 2 hours at 37°C. Inoculated cells were loaded with CCF2-AM at 15°C for 1 hour, incubated overnight at room temperature, and then analyzed by flow cytometry. Quantification of the relative percentages of cleaved CCF2^+^ cells (fusion efficiency) in total cells are shown. Data are presented as mean ±SD. The experiments were performed three times with PBMCs (A) and primary CD4^+^ T cells (B) isolated from three donors. The *p* values were determined using two-tailed paired Student′s *t*-test. ***, *p*<0.001; n.s., not significant.

**Figure S8.** Validation of labelling efficiency of HIV-1_NL4-3/Gag-iVenus_ with mScarlet-Vpr and the infectivity of HIV-1_NL4-3/Gag-iVenus_ containing mScarlet-Vpr and HIV-1_NL4-3/MA-3xHA_. (A) HIV-1_NL4-3_, HIV-1_NL4-3/Gag-iVenus_, and HIV-1_NL4-3/Gag-iVenus_ containing mScarlet-Vpr were plated on top of poly-L-lysin-coated glass bottom chambers, and fixed. The virus particles were observed with the spinning-disk confocal microscope. The fraction of mScarlet-positive particles relative to total Venus-positive particles is shown in the right panel. More than 8000 virus particles were analyzed. Bars, 10 μm. (B) The infectivity of HIV-1_NL4-3,_ HIV-1_NL4-3/Gag-iVenus_ containing mScarlet-Vpr, and HIV-1_NL4-3/MA-3xHA_ was measured by TZM-bl assay. Data are presented as mean ±SD. Three independent experiments were performed. The *p* values were determined using Dunnett’s test following one-way ANOVA. *, *p*<0.05; ***, *p*<0.001

**Figure S9.** Unmodified HIV-1 can be internalized into activated primary CD4+ T cells through macropinocytosis. Activated primary CD4^+^ T cells were inoculated with HIV-1_NL4-3_ labeled with DiOC18 and R18 in the presence of BSA-AlexaFluor647. Cells were labeled with an anti-CD4 antibody conjugated with BV421, plated on top of a poly-L-lysin-coated glass bottom chamber, and fixed. The fixed cells were observed with a spinning-disk confocal microscope. Individual *z*-slices of representative cells are shown. Yellow arrowheads indicate the R18 signal without the DiOC18 signal colocalized with BSA-AlexaFluor647 (HIV-1 internalization through macropinocytosis). White arrowheads indicate the dequenching of DiOC18 in the presence and absence of a fusion inhibitor AMD3100. Bar, 10 μm.The experiments were performed two times with primary CD4^+^ T cells isolated from two donors.

**Figure S10.** Insertion of MA-3xHA into the stalk region of MA does not affect virus release, and maturation. Western blot analysis for the release and maturation of HIV-1_NL4-3_, HIV-1_NL4-3/MA-3xHA,_ produced from 293T cells. Representative western blotting images from one out of three independent experiments were shown.

**Figure S11.** Schematic illustration of the procedure to classify MA-3xHA signals into three groups, a) not associated with macropinosomes, b) associated with permeabilized macropinosomes, and c) associated with intact macropinosomes.

**Figure S12.** J/B inhibits HIV-1 infection of primary CD4^+^ T cells. Activated primary CD4^+^ T cells were pretreated with either J/B or vehicles for 15 minutes at 37°C and inoculated with HIV-1_NL-NI_ for 2 hours at 37°C. Cells were washed and cultured for 24 hours at 37°C in the presence of saquinavir. Cells were then lysed, and the NanoLuc activity in the cell lysates was measured. The NanoLuc activity was normalized by total protein amounts. Data are presented as mean +SD. The experiments were performed with primary CD4^+^ T cells isolated from three donors in three independent experiments. The *p* values were determined using two-tailed paired Student′s *t*-test. **, *p*<0.01.

## Notes

### Competing Interest Statement

The authors have declared no competing interest.

