## Supplementary figures and images for "Macropinosomes are a site of HIV-1 entry into primary CD4^+^ T cells"

### Figure S1

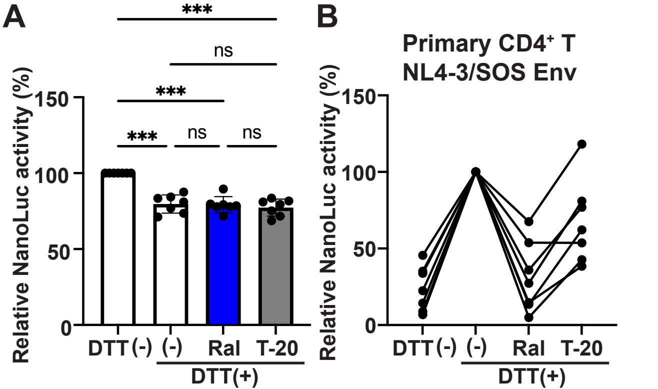

### Figure S2

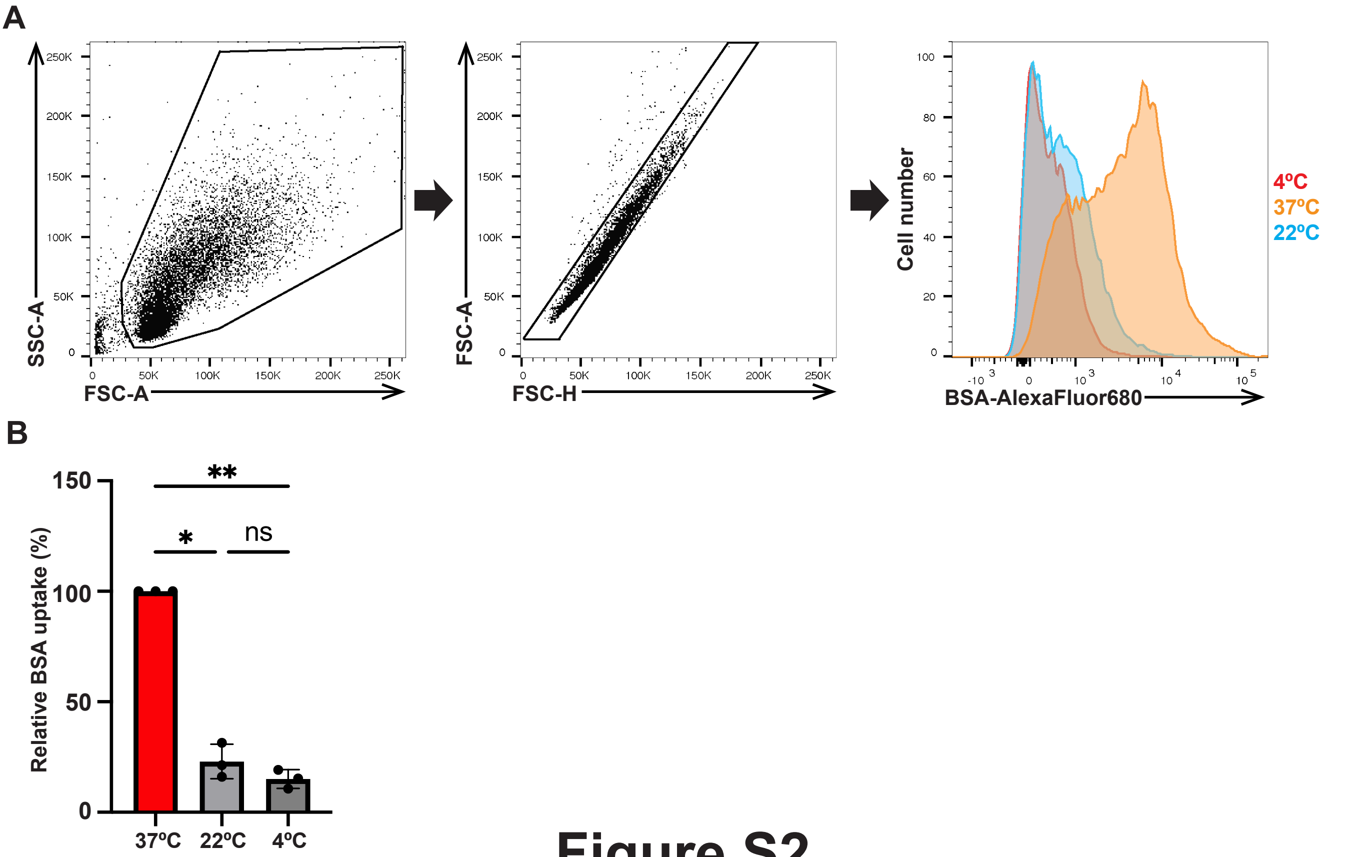

### Figure S3

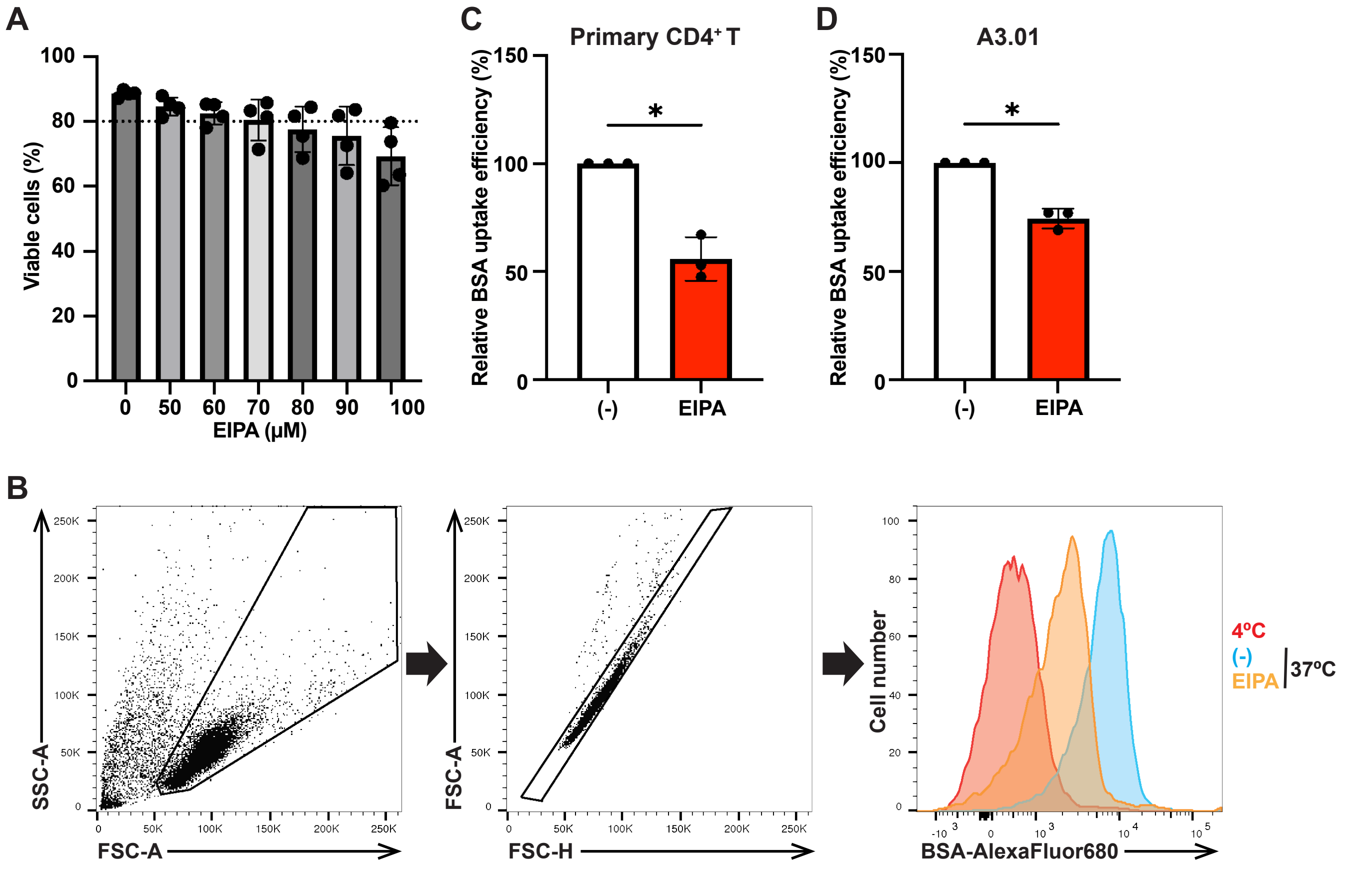

### Figure S4

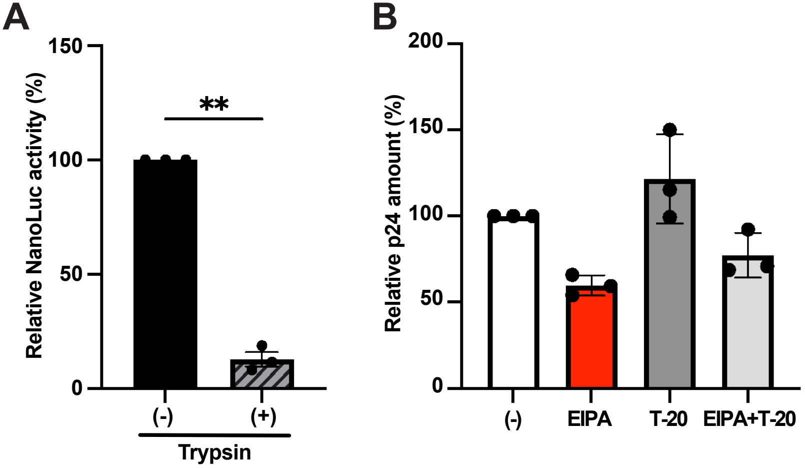

### Figure S5

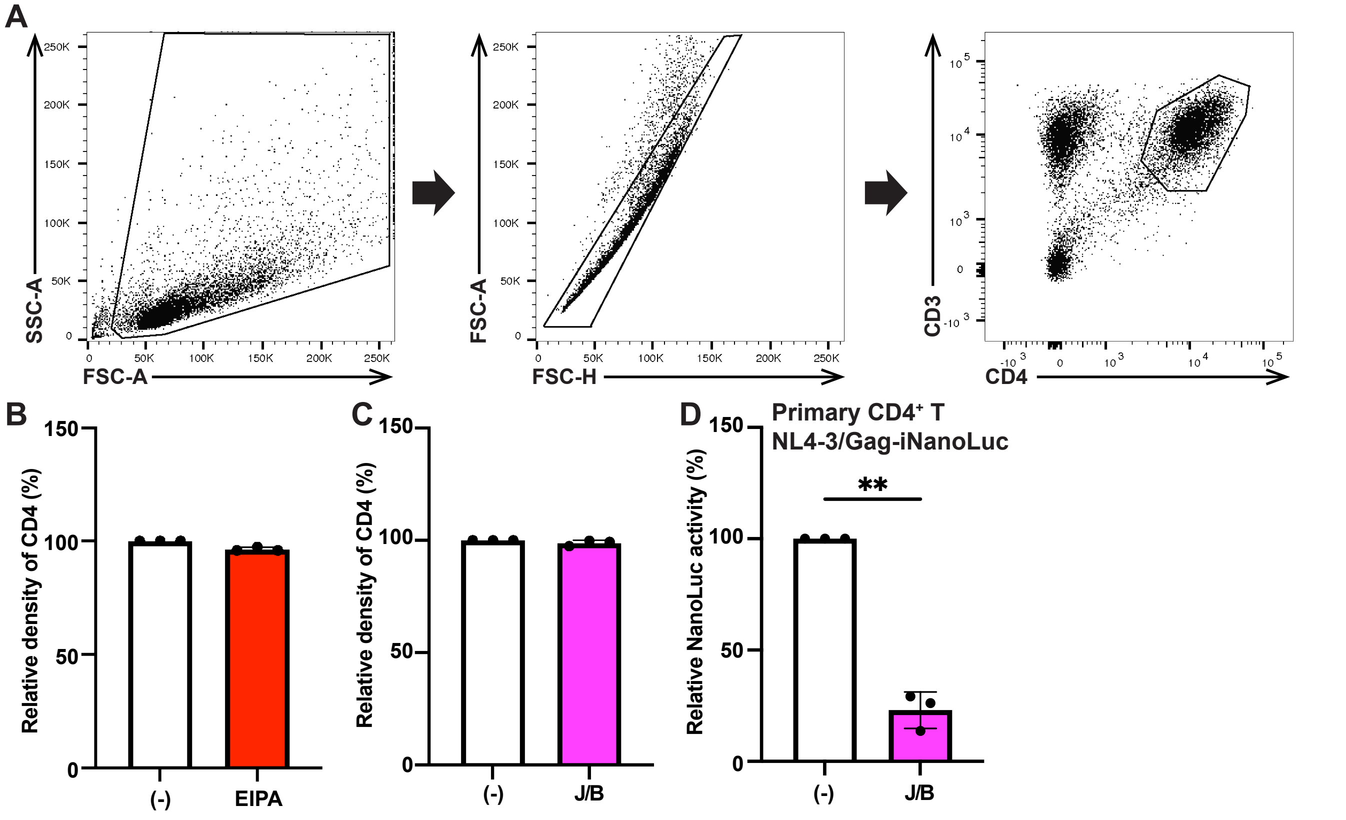

### Figure S6

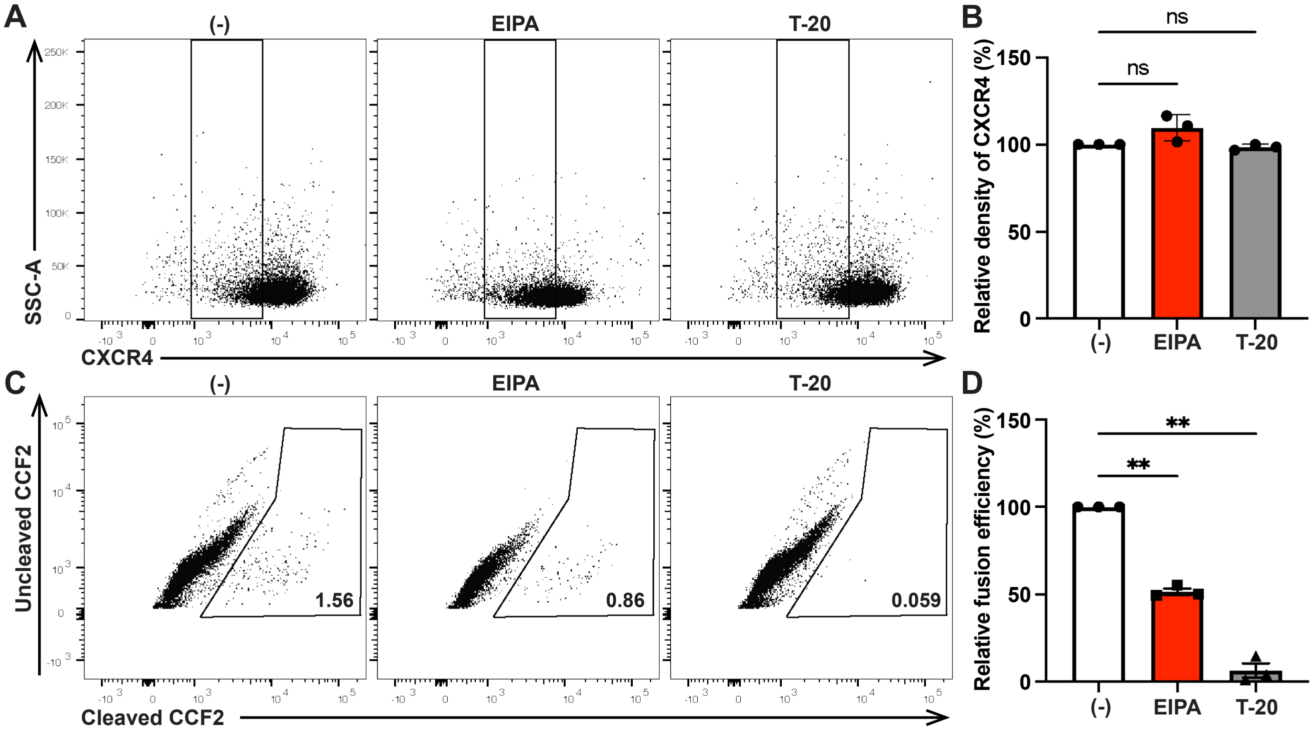

### Figure S7

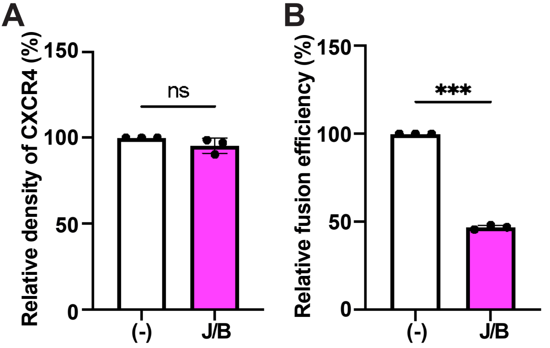

### Figure S8

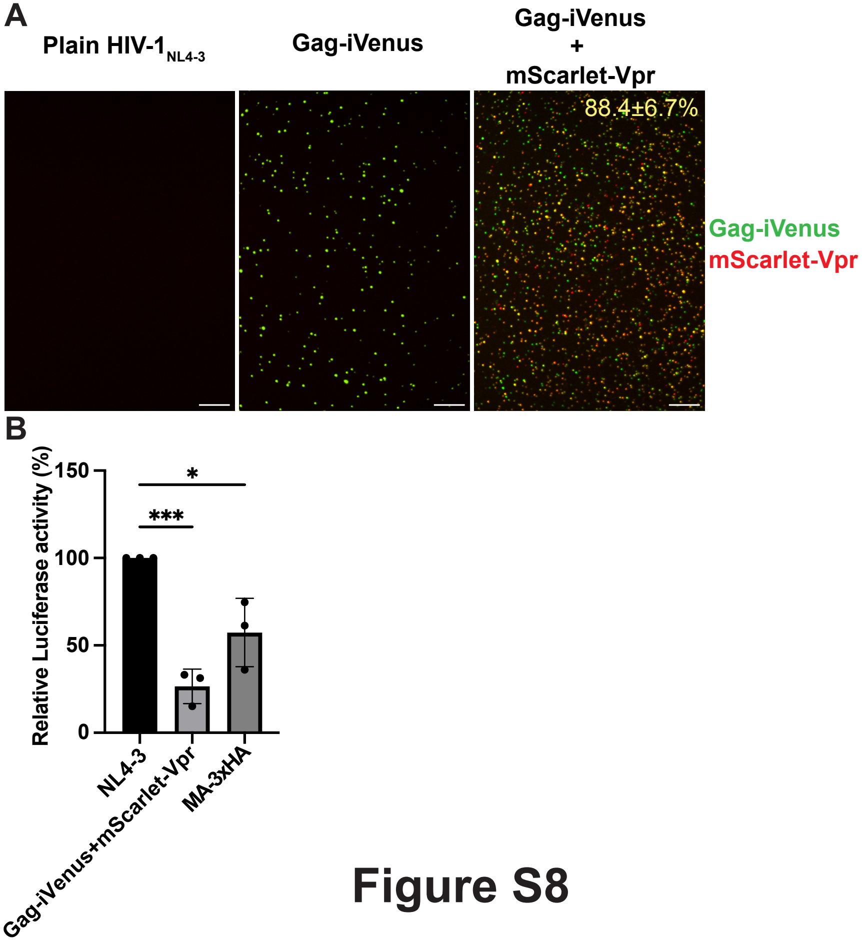

### Figure S9

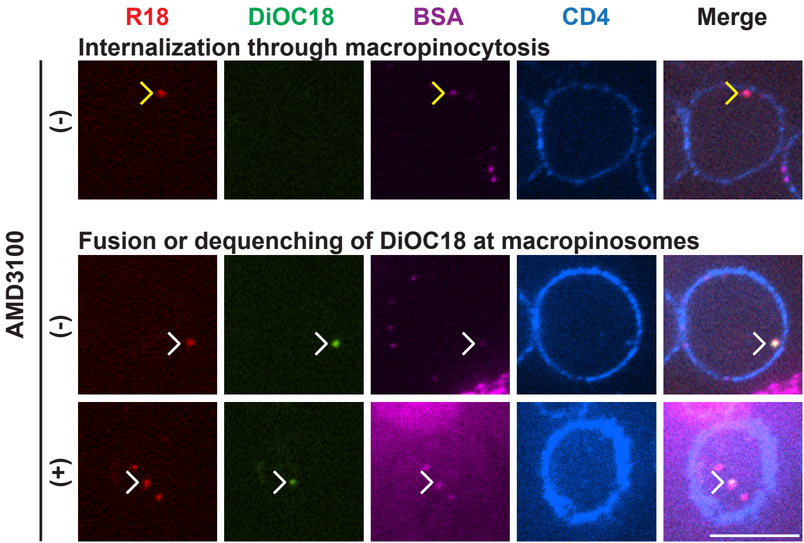

### Figure S10

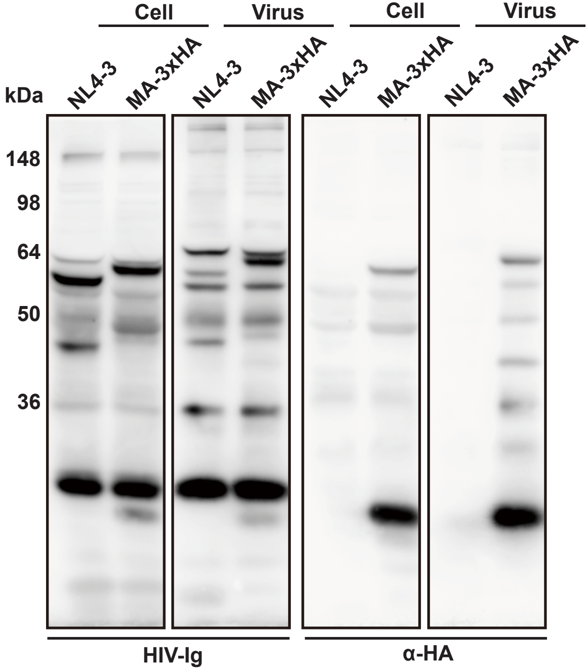

### Figure S12

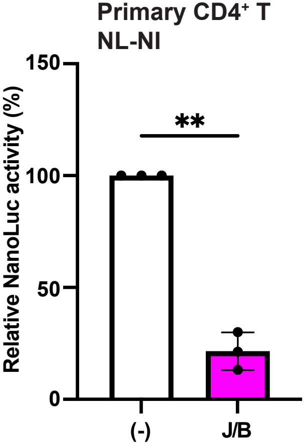


#### Figure S12
