## Supplementary material for "Macropinosomes are a site of HIV-1 entry into primary CD4^+^ T cells": Figure S11

1) Measure the radii of each BSA-AlexaFluor488 spot and determine $R_{488}$
Ninety-five percentages of BSA-AlexaFluor488 spots have the same radii as or shorter radii than $R_{488}$

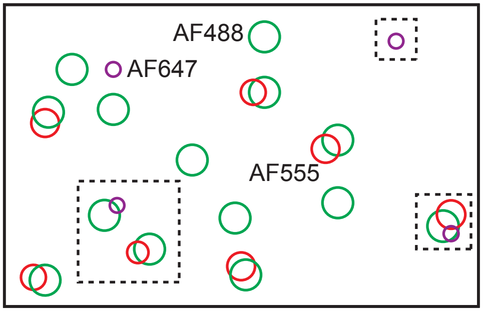


2) Measure the distances from each BSA-AlexaFluor488 spot to the nearest AlexaFluor555 spot (Shortest Distance488-555; $SD_{488\text{-}555}$) and to the nearest AlexaFluor647 spot (Shortest Distance488-647; $SD_{488\text{-}647}$), respectively

3) Sort MA-3xHA spots into three classifications
(a’) $R488 \geqq SD_{488\text{-}555}$ and $R488 \geqq SD_{488\text{-}647}$
(b’) $R488 < SD_{488\text{-}555}$ and $R488 \geqq SD_{488\text{-}647}$
(c’) Neither (a’) nor (b’)

4) Validate the classifications by comparing the radius of each BSA-AlexaFluor488 spot with $SD_{488\text{-}647}$ and $SD_{488\text{-}555}$
(a) MA-3xHA associated with permeabilized macropinosome; $r \geqq SD_{488\text{-}555}$ and $r \geqq SD_{488\text{-}647}$
(b) MA-3xHA associated with Intact macropinosome; $r < SD_{488\text{-}555}$ and $r \geqq SD_{488\text{-}647}$
(c) MA-3xHA not associated with macropinosome; neither (a) nor (b)

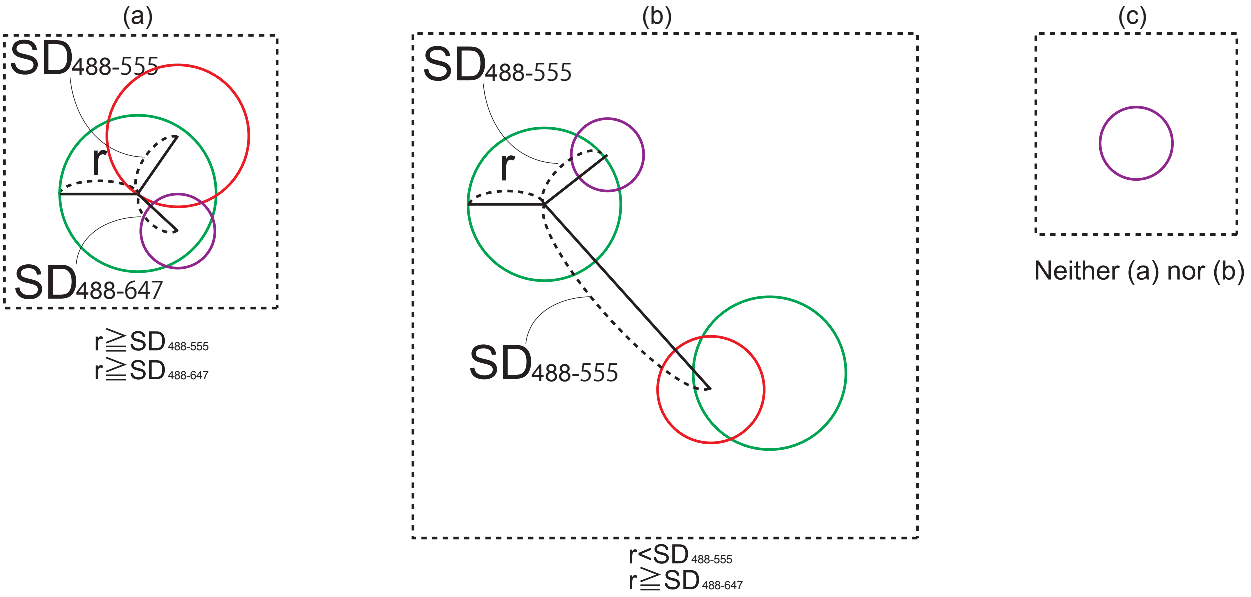


### Figure S11
